# Internal modifications in the CENP-A nucleosome modulate centromeric dynamics

**DOI:** 10.1101/101006

**Authors:** Minh Bui, Mary Pitman, Arthur Nuccio, Serene Roque, Paul Gregory Donlin-Asp, Aleksandra Nita-Lazar, Garegin A. Papoian, Yamini Dalal

## Abstract

Post-translational modifications (PTMs) of core histones have studied for over 2 decades, and are correlated with changes in transcriptional status, chromatin fiber folding, and nucleosome dynamics. However, within the centromere-specific histone H3 variant CENP-A, few modifications have been reported, and their functions remain largely unexplored. In this multidisciplinary report, we utilize *in silico* computational and *in vivo* approaches to dissect lysine 124 of human CENP-A, which was previously reported to be acetylated in advance of replication. Computational modeling demonstrates that acetylation of K124 causes tightening of the histone core, and hinders accessibility to its C-terminus, which in turn diminishes CENP-C binding. Additionally, CENP-A K124ac/H4 K79ac containing nucleosomes are prone to DNA sliding. *In vivo* experiments using an acetyl or unacetylatable mimic (CENP-A K124Q and K124A respectively) reveal alterations in CENP-C levels, and a modest increase in mitotic errors. Furthermore, mutation of K124 results in alterations in centromeric replication timing, with the permanently acetylated form replicating centromeres early, and the unacetylable form replicating centromeres late. Purification of native CENP-A proteins followed by mass spectrometry analysis reveal that while CENP-A K124 is acetylated at G1/S, it switches to monomethylation during early and mid-S phase. Finally, we provide evidence that the HAT p300 is involved in this cycle. Taken together, our data suggest that cyclical modifications within the CENP-A nucleosome can influence the binding of key kinetochore proteins, the integrity of mitosis and centromeric replication. These data support the emerging paradigm that core modifications in histone variant nucleosomes transduce defined changes to key biological processes.

## INTRODUCTION

Post-translational modifications in histones play an important role in chromosome biology. The majority of such modifications discovered exist on the N-terminal tails of histones H3, H2A, H3.3 and H4 (Davie and Candido, 1978; Rando, 2012; Rothbart and Strahl, 2014). N-terminal histone modifications may increase nucleosome turnover (Ferrari and Strubin, 2015), be inherited at specific loci (Xie *et al.,* 2015), alter the binding efficiency of various transcriptionally active or repressive factors (Tropberger *et al.,* 2013), and disrupt replication timing (Pryde *et al.*, 2009), thereby potentially influencing the fate of the underlying locus. A new area of research has also uncovered covalent modifications within histone fold domains, such as H3K56ac and H3K122ac (Maas *et al.*, 2006; Schneider *et al.*, 2006; Manohar *et al.*, 2009). H3K122 is acetylated at the nucleosome dyad, wherein it alters DNA-histone binding and increases thermal repositioning of the nucleosome *in vitro* (Manohar *et al.*, 2009). Concurrently, *in vivo* experiments demonstrate that H3K122ac promotes nucleosome turnover, thereby stimulating transcription (Hainer and Martens, 2011; Devaiah *et al.*, 2016). A single residue in the histone variant H3.5, leucine 103, when mutated disrupts nucleosome instability both *in vitro* and *in vivo* (Urahama *et al.*, 2016). Indeed, a single change in the nucleosome, methylation at H3K9, alters replication timing by modulating the binding of ORC2 (Giri *et al.*, 2015). Thus, internal histone core modifications or key residue alterations can alter the nucleosome structure in a manner distinct from that reported for histone tail modifications. (Taverna *et al.*, 2007; Molina-Serrano and Kirmizis, 2013; Tropberger *et al.*, 2013). Therefore, investigating how histone fold domain modifications contribute to structure or function, especially in histone variants which mark specific loci, such as the centromere-specific histone variant CENP-A, is an exciting area of research.

A plethora of CENP-A modifications have been discovered in recent years (Bailey *et al.*, 2013); however, only 3 have been reported within the histone fold domain (Bui *et al.*, 2012; Niikura *et al.*, 2015; Yu *et al.*, 2015; Niikura *et al.*, 2016). Using exogenously and constitutively expressed epitope tagged CENP-A, mass spectrometric analysis uncovered phosphorylation of S68 within loop 1 (Yu *et al.*, 2015), and ubiquitination at K124 near the pseudo-dyad (Niikura *et al.*, 2015; Niikura *et al.*, 2016). In addition, analysis of chromatin bound native CENP-A (nCENP-A) identified acetylation of K124 (K124ac) specifically at G1/S phase (Bui *et al.*, 2012). K124 in CENP-A is analogous in location to residue K122 in histone H3, which, as discussed above, has a significant impact on nucleosome structure and function. We previously reported that in advance of replication, CENP-A K124ac and H4K79ac correlates with a transitionary state of native CENP-A nucleosomes, and with the opening of centromeric chromatin fiber (Bui et al 2012). Inheritance of nucleosomes post-replication involves a coordinated and complex series of events that include eviction of nucleosomes, followed by the rapid reassembly of disrupted nucleosomes after the passage of the replication machinery (Ramachandran and Henikoff, 2015). CENP-A K124ac is proximal to both, the pseudodyad DNA of the CENP-A nucleosome, and to the CENP-A C-terminus, which is required to bind the inner kinetochore protein CENP-C (Carroll *et al.*, 2010). Therefore, we hypothesized that potential functions of CENP-A K124ac coupled to H4K79ac might be to alter the stability of the CENP-A nucleosome, or, to alter the binding of the kinetochore protein CENP-C. Cumulatively, we suggested, such events might be necessary to increase access to the centromeric chromatin fiber at the appropriate time in replication. Consequently, constitutive gain or loss of this G1/S specific modification in CENP-A might be expected to influence centromere replication dynamics.

In this report, we dissect the role of CENP-A K124ac *in silico* and *in vivo*. First, using all-atom computational modeling, we simulate the presence of K124ac and H4K79ac in the octameric CENP-A nucleosome, finding that it results in a loosening of DNA at the pseudo-dyad, followed by asymmetric site exposure of the terminal ends of the DNA. These modification-induced DNA dynamics are driven, in part, by an unexpected compaction of the CENP-A protein, accompanied by a locking of the CENP-A C-terminus. Consistent with this finding, further computational analysis of CENP-A nucleosomes, which contain CENP-A K124ac and H4K79ac, show dramatically reduced contacts with CENP-C. To examine the function of CENP-A K124ac *in vivo*, we generated mutants of CENP-A K124 which mimic either the constitutively acetylated (K124Q) or the unacetylated state (K124A), and find three aspects of centromere dynamics are impacted. First, gain or loss of CENP-A K124ac results in a quantifiable decrease in association of modified CENP-A nucleosomes to the kinetochore protein CENP-C. Second, there is a modest accumulation of downstream mitotic errors. Third, relative to wildtype CENP-A, in cells expressing the un-acetylatable CENP-A K124A, centromeric foci are delayed in replication timing, whereas in cells expressing CENP-A K124Q, centromeric foci speed up through replication. Using purification and subsequent mass spectrometry analysis of native CENP-A at distinct points spanning G1/S to late S phase, we uncover a switch in modifications of chromatin-bound native CENP-A, from acetylation of K124 at G1/S to monomethylation of K124 at S phase. Finally, using two custom antibodies against CENP-A K124ac, our data suggest the involvement of p300 as a putative HAT which appears to promote acetylation of chromatin-bound native CENP-A.

Together, these results suggest a working model wherein cyclical control of CENP-A K124 modifications may serve as binary toggle for modulating association of CENP-A to CENP-C, thereby assisting in facilitating accurate timing of centromeric replication and to mitotic integrity.

## RESULTS

In recent work, using all-atom molecular dynamics (MD), we showed that the 4-helix bundle in the CENP-A nucleosome core particle (CENP-A NCP), (**Fig 1A**) is highly dynamic (Winogradoff *et al*., 2015b), such that CENP-A dimers were found to move antiparallel to each other in a “shearing” motion unique to octameric CENP-A nucleosomes. Earlier experimental work from our lab had indicated that G1/S CENP-A nucleosomes are enriched in CENP-A K124ac and H4K79ac, and occupy a transitionary form, whereas the majority of CENP-A nucleosomes are stable octamers in early S phase (Bui *et al*., 2012). To investigate potential transitionary dynamics, we computationally modeled CENP-A K124ac and H4K79ac in the context of the octameric CENP-A nucleosome. Four systems were simulated: (1) the CENP-A NCP and (2) the acetyl CENP-A NCP with four acetylations –K124ac of CENP-A and CENP-A’ and K79ac of H4 and H4’, (3) the same two species above, bound to a fragment of the kinetochore protein CENP-C.

**Figure 1.**
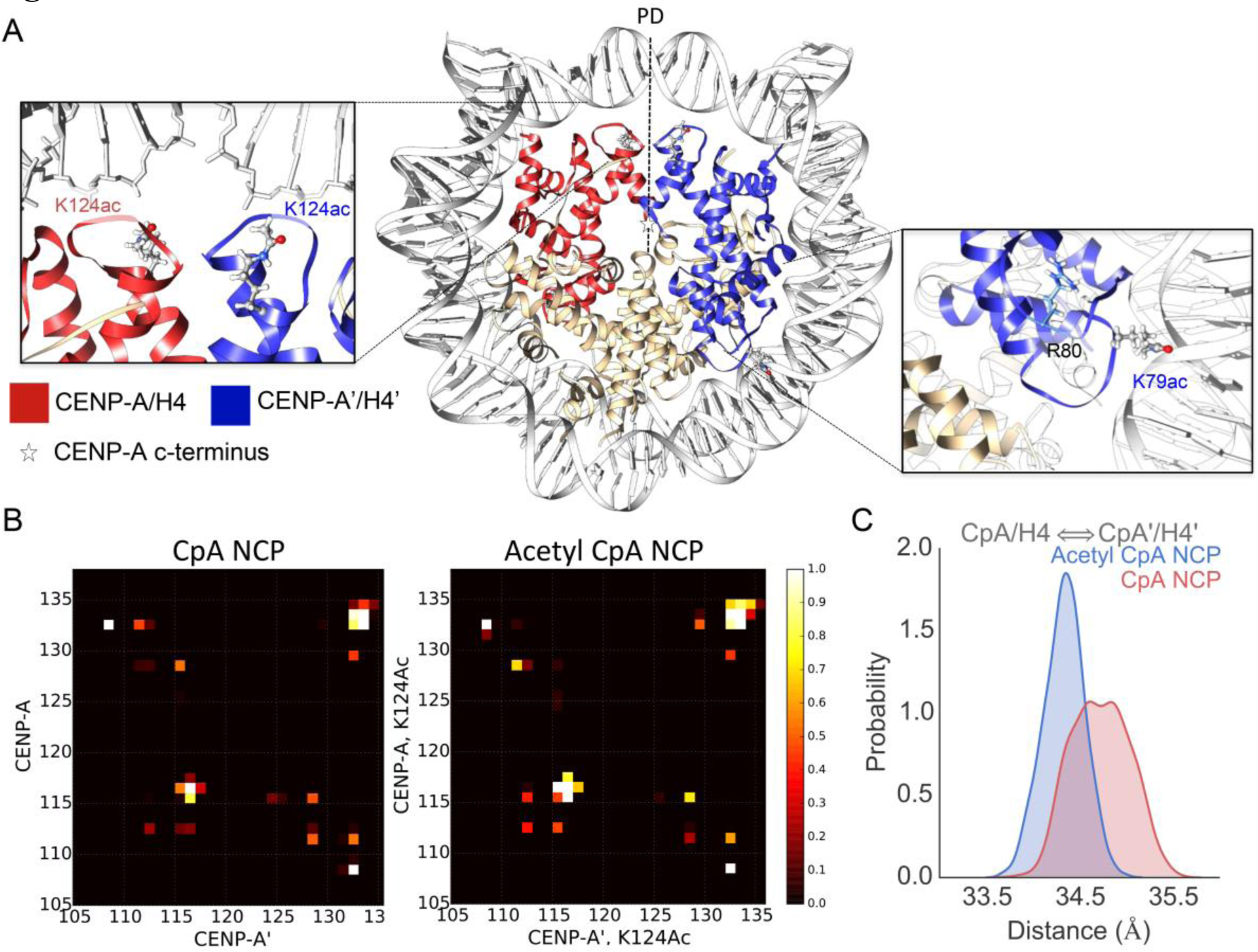
The CENP-A nucleosome displays a tightening of the histone core upon lysine acetylation. **A)**The starting structure of the acetyl CENP-A NCP is shown with CENP-A and H4 in red, CENP-A’ and H4’ in blue, and H2A/H2B monomers in tan. The pseudo-dyad is shown by a dotted black line and marked as PD. Overlay’s shown K124ac, K79ac, and CENP-A R80 and the L1 loop in more detail. This structure is after 1 μs simulation of CENP-A and the production of the starting structure is described (Methods). **B)** Contact Analysis showing CENP-A (CpA) to CENP-A’(CpA’) interface at the 4-helix bundle. The contact cutoff between Cα atoms was set to 8Å. An increase of contacts is shown upon acetylation—a decrease in the 4-helix bundle interface distance. A value of 1, white, shows a contact formed over all simulation time steps and 0, black, never. In the acetyl NCP residues H115, A116, and G117 more frequently form contacts located in the hinge region of the 4-helix bundle. The C-terminus, specifically CENP-A’ residues I132, R133, and G134, forms increased contacts with CENP-A G134. **C)** The center-of-mass (COM) of dimers was measured over all time steps and the resulting distributions are shown for CENP-A/H4 to CENP-A’/H4’, the acetylated histones. The acetylated system, shown in blue, shows a decrease in variance and distance between the two dimer COMs.

## The acetylated CENP-A nucleosome displays a tightening of the histone core

Due to the loss of positive charge on CENP-A K124ac and H4 K79ac, we postulated that the association of the histone core with DNA and inter-histone repulsion would both decrease. We tested this hypothesis with detailed contact analysis and found that the acetyl NCP 4-helix bundle interface makes a greater proportion of contacts throughout the simulation (**Fig 1A-B**). In the acetyl NCP residues H115, A116, and G117 more frequently form contacts located in the hinge region of the 4-helix bundle. The constraint on this flexible hinge stabilizes the CENP-A to CENP-A’ interface. As a result, the C-terminus –specifically CENP-A’ I132, R133, and G134– forms more contacts with CENP-A G134 (**Fig 1B**).

To explore these dynamics in greater detail, we next performed Principle Component Analysis of histone core proteins (PCA^core^). This method dissects small local vibrations out to clarify large global distortions (PC modes). In this analysis, the most significant mode of motion, PC1^core^, revealed a surprising “freezing” or lack of motion at histone interfaces at the pseudo-dyad, adjacent to the K124ac modifications (**Fig 2** and **Movie S1**). Because of the decreased histone rocking in the acetyl NCP, even histone interfaces far away from the studied modifications compact, hence, the effect of acetylated lysines transduce to the other face of the nucleosome (**Fig 3A** **and Movie S1**).

**Figure 2.**
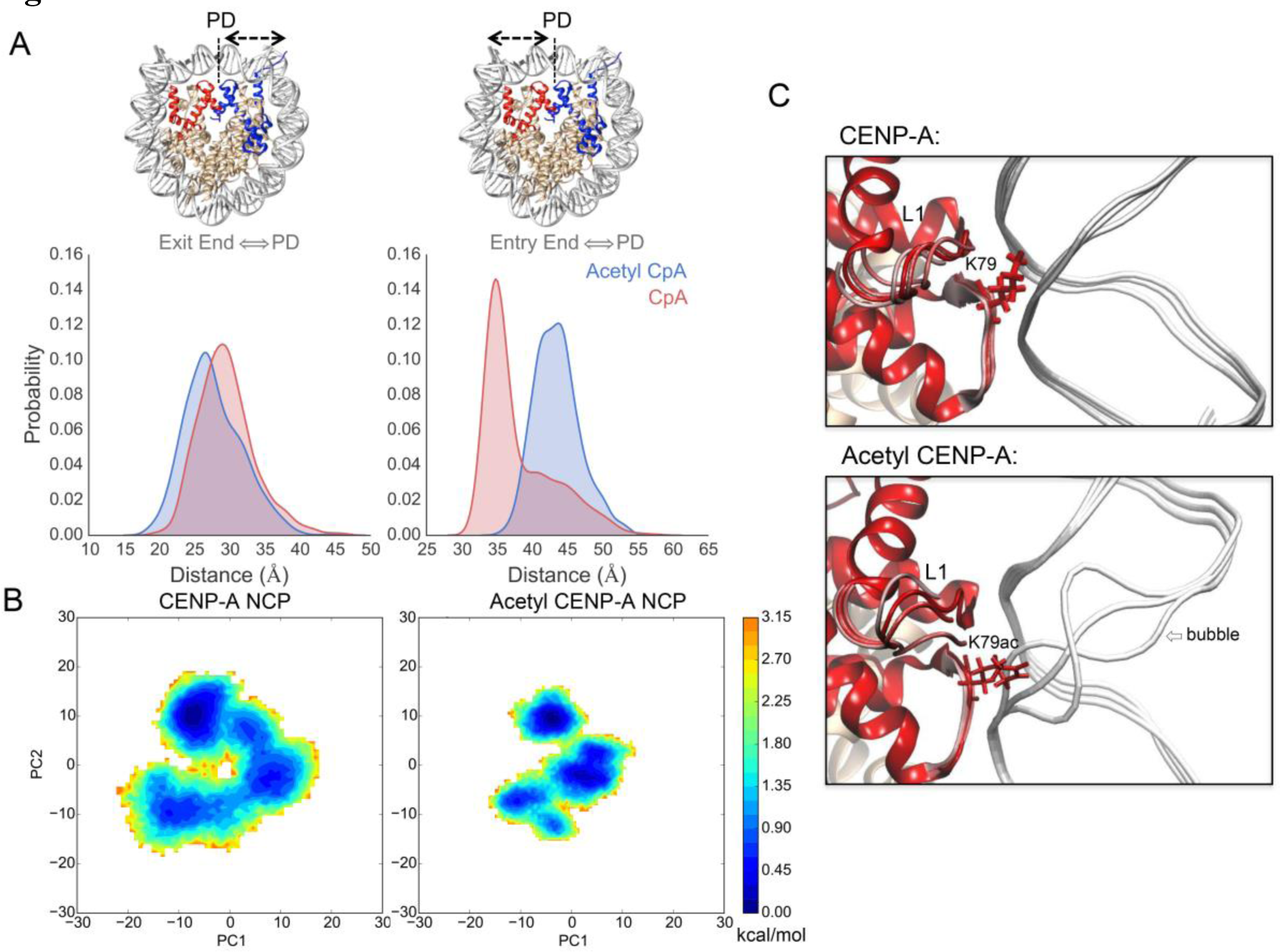
Lysine Acetylations asymmetrically loosen DNA entry and exit ends and alter DNA dynamics. **A)** The distance between DNA ends to the pseudo-dyad was measured for all time steps and distributions shown. In the structures showing which graph corresponds to which DNA end, the CENP-A NCP is shown with CENP-A and H4 in red, CENP-A’ and H4’ in blue, and H2A/H2B monomers in tan. The entry end of DNA unwraps more in the acetyl NCP in blue. **B)** Coarse-grained free energy landscapes are shown for both systems through the projection of PC1^NUC^ and PC2^NUC^ of the whole nucleosomal Principal Component Analysis (PCA^NUC^). Here it is shown that the Acetyl NCP landscape becomes more rugged and frustrated. This means that the system is less freely sampling a larger phase space as seen through the attenuation of histone rocking in supplemental movies. **(C)** From PC1^NUC^ trajectories, three representative snap shots are shown to illustrate the intra-helical bubble formed in DNA near K79ac that does not occur in unmodified CENP-A. DNA is shown in white, and the L1 loop of CENP-A is marked.

**Figure 3.**
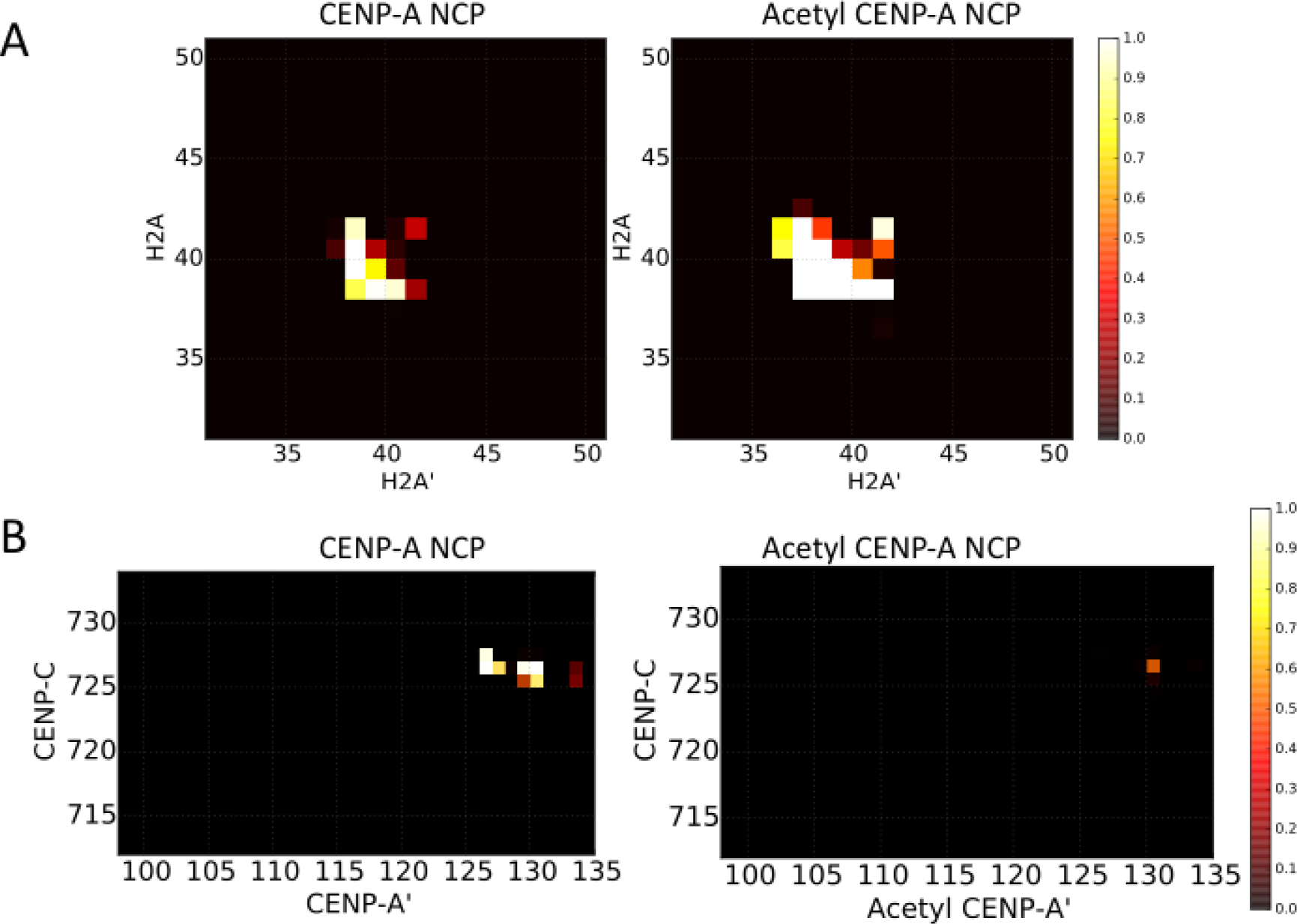
The global compaction of the histone core disrupts CENP-C binding. **A)** Contact Analysis of the H2A to H2A’ interface on the opposite face of the nucleosome from the acetylations studied. The contact cutoff between Cα atoms was set to 8 Å. A value of 1, white, shows a contact formed over all simulation time steps and 0, black, never. An increase of contacts is shown upon acetylation consistent with a global compaction of the histone core transduced away from the points of acetylation. **B)** To study the effects of the histone core compaction with acetylation, the CENP-C fragment (Kato *et al*. 2013) was docked to the nucleosomes and contacts analyzed. The CENP-C fragment is shown to form stable contacts with the CENP-A c-terminal end that are lost upon acetylation.

We next wished to assess whether only the histone interfaces tighten in the acetyl CENP-A NCP or whether the globular histone domains also contract. To investigate this, we calculated the distance between the center-of-mass (COM) of dimers. We found that the dimers CENP-A/H4 and CENP-A’/H4’ are closer together in the acetyl NCP (**Fig 1C**).

In other dimer combinations, such as CENP-A/H4 to H2A/H2B, the variance in the distance between dimers decreased, consistent with the rigidification of the histone core upon acetylation. Overall, the observed changes in distances between various dimers show that the histone core is stabilized and tightened in the acetyl NCP.

To compare local structural flexibility, we calculated the root-mean-square fluctuation (RMSF) of Cα atoms (**Fig S1**). Compared to the control, the RMSF of the acetyl CENP-A NCP was locally suppressed in specific regions, namely, the C-terminus of CENP-A and the acidic patch of H2A. Both of these regions are targets for CENP-C binding (Kato *et al*., 2013). A diminution in the accessibility of the acidic patch may interfere with H2A D89 binding with CENP-C R717 and R719 (Kato *et al*., 2007). This unexpected and distinct reduction in the availability of the C-terminus of CENP-A and H2A acidic patch emphasizes local changes that result from these acetylations. In order to test if acetylation modulates CENP-C docking, the CENP-C fragment from the recently solved co-crystal of H3:CENP-A plus CENP-C (Kato et al., 2015) was docked to each system and simulated for an additional microsecond. As can be seen, the CENP-C fragment forms stable contacts with the C-terminus of CENP-A in the unmodified state, which are virtually lost upon acetylation (**Fig 3B**). The acetylations modeled result in global structural changes that are sufficient to diminish the accessibility of a critical CENP-C docking interface. Interestingly, in the acetyl NCP, the heterotetramer half showing the greatest suppression of RMSF is also the region with increased DNA unwrapping (**Fig 2A** and **Fig S1**). To further study the interaction of histones and DNA, we then extended our analysis to the whole nucleosome.

## Specific Lysine Acetylations increase accessibility of DNA in acetyl NCP

In experiments, we observed that the DNA in both systems, CENP-A and CENP-A K124ac asymmetrically unwraps, but has enhanced unwrapping in the acetyl NCP (**Fig 2A** and **S1**). We studied the change in DNA dynamics further with Principle Component Analysis of the whole nucleosome (PCA^NUC^). Visual analysis of the most significant mode, PC1^NUC^, demonstrated a pronounced untwisting motion of DNA ends in the acetyl NCP (**Movie S2**).

Another feature exclusive to PC1^NUC^ of the acetyl NCP is a modulation in the widths of the major and minor grooves of DNA (**Movie S2**). The cause for this modulation was a pronounced scissoring motion between helices α2 and α3 of the 4-helix bundle in acetyl CENP-A. We observed a high coherence between the scissoring of the 4-helix bundle with the modulation of the size of the DNA minor grooves (correlation coefficient of 0.82, methods). This suggests that the altered motion of the 4-helix bundle in acetyl CENP-A could promote DNA sliding. PCA^NUC^ also revealed that adjacent to both H4 and H4’ K79ac, two intermittent DNA bubbles formed within the double helix (**Movie S2**) indicating that these regions of DNA become more susceptible to opening in the presence of H4K79ac.

Overall, these all-atom computational modeling results lead to two discrete possibilities. First, the rigidifying of the acetyl NCP in comparison to CENP-A NCP locks the CENP-A C-terminus and stabilizes the H2A acidic patch, which could potentially disrupt protein access to the CENP-A C-terminus; and second, asymmetric exposure of the DNA coupled with DNA bubbling may promote DNA sliding and thereby open the centromeric chromatin fiber to prime centromeric replication. To investigate these possibilities further, we turned to an experimental approach *in vivo*.

## CENP-A K124 acetylation is not required for accurate targeting to centromeres

To dissect the function CENP-A K124 modifications *in vivo*, we used site directed mutagenesis to generate versions of CENP-A K124 which mimic either a constitutively un-acetylated state (Lysine to Alanine, K124A); or, a constitutively acetylated state (Lysine to Glutamine, K124Q). These mutants were used for the series of experimental investigations discussed below (scheme in **Fig S2A**).

We observed virtually no defects in centromeric localization for these mutants, with almost complete centromeric co-localization with mCh-CENP-A (**Fig 4A**). To confirm this result, we co-immuno-stained K124A/Q along with CENP-C, and found that K124A/Q foci co-localized with CENP-C foci (**Fig 4B**). Furthermore, when K124A or K124Q were immuno-stained with other centromeric markers, including CENP-B, these mutant foci also co-localized with the centromeric markers in a vast majority of the cells, throughout the cell cycle (**Fig 4**, **S3A**, **S3C**, **6** and **7**). Although we did not dissect the efficiency of assembly of mutants relative to wildtype CENP-A, taken together these results suggest that both CENP-A K124A and K124Q assemble at centromeres in a manner visually comparable to wildtype CENP-A.

**Figure 4.**
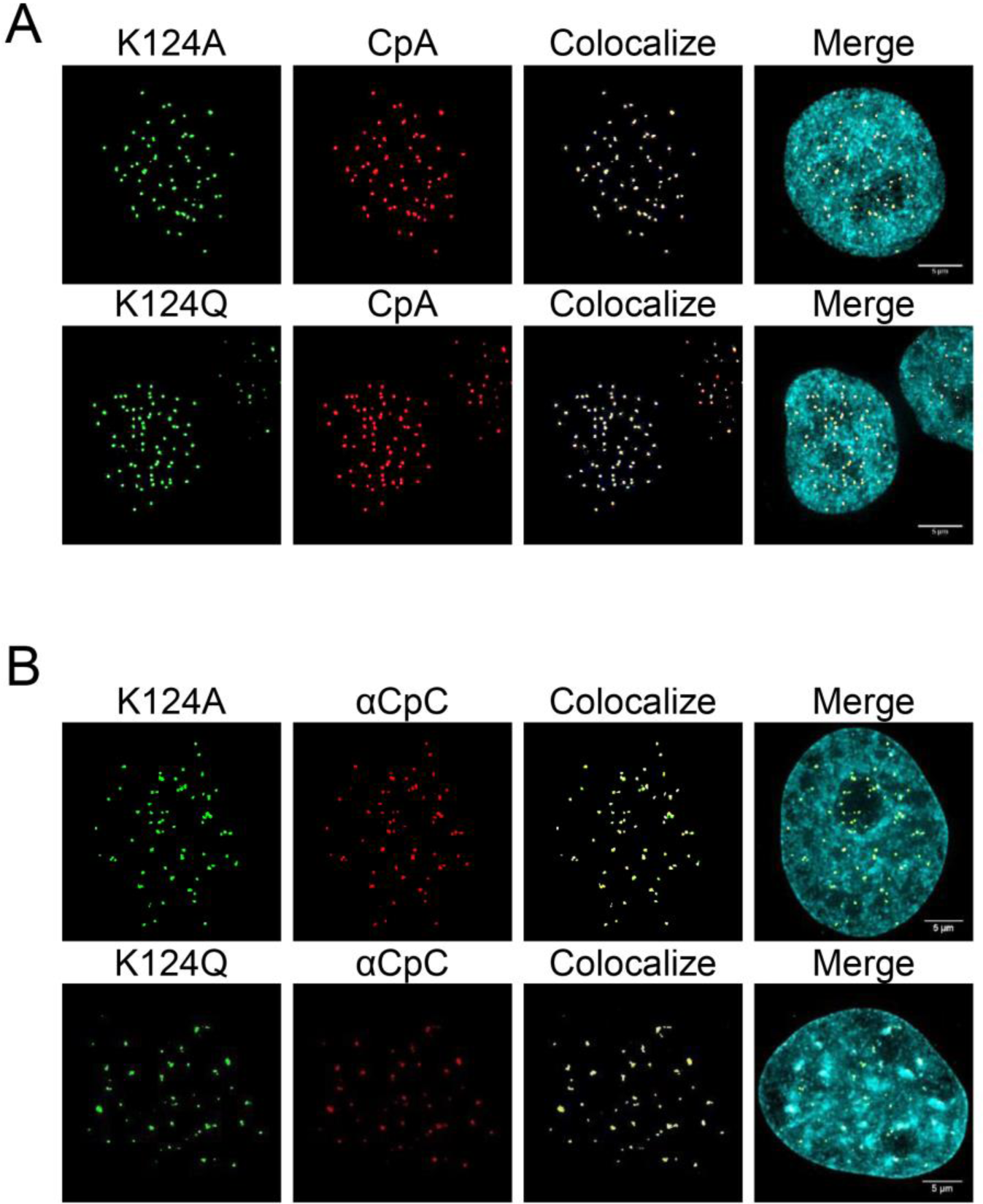
K124A/Q display normal centromeric localization. **A)** CENP-A K124A/Q were fused to an N-terminal GFP tag and co-expressed with mCh-CENP-A (CpA). The colocalization finder plug-in in ImageJ was used to find colocalizing GFP- and mCh- foci to indicate centromeric deposition, and final panel depicts a complete merge of all three channels including DAPI. **B)** CENP-C (CpC) immuno-fluorescence (IF) was performed on GFP-tagged CENP-A K124A/Q to look for colocalization.

## Constitutive gain or loss of K124 acetylation results in alterations in CENP-C binding

One key computational prediction from *in silico* experiments above (**Fig 1**–**3**) was that the acetyl CENP-A nucleosomes have reduced accessibility at the C-terminus, potentially impacting protein binding to this domain, most plausibly CENP-C, which binds the C-terminus of CENP-A (Carroll *et al*., 2010; Kato *et al*., 2013). Therefore, we tested whether K124 acetylation impacts the interaction between modified CENP-As and CENP-C *in vivo*. We fused CENP-A, K124A, and K124Q to an N-terminal epitope HA-tag. This strategy avoids interference with the recognition motif for CENP-C on the CENP-A C-terminal tail (Carroll *et al*., 2010; Kato *et al*., 2013).

We first confirmed whether these mutant CENP-A proteins were expressed normally in whole cell extracts (WCE). WCE for total protein levels revealed that the exogenously expressed HA-tagged proteins, the CENP-A chaperone HJURP, and the inner kinetochore protein CENP-C levels remained relatively equal in all three cell lines (**Fig 5A**).

**Figure 5.**
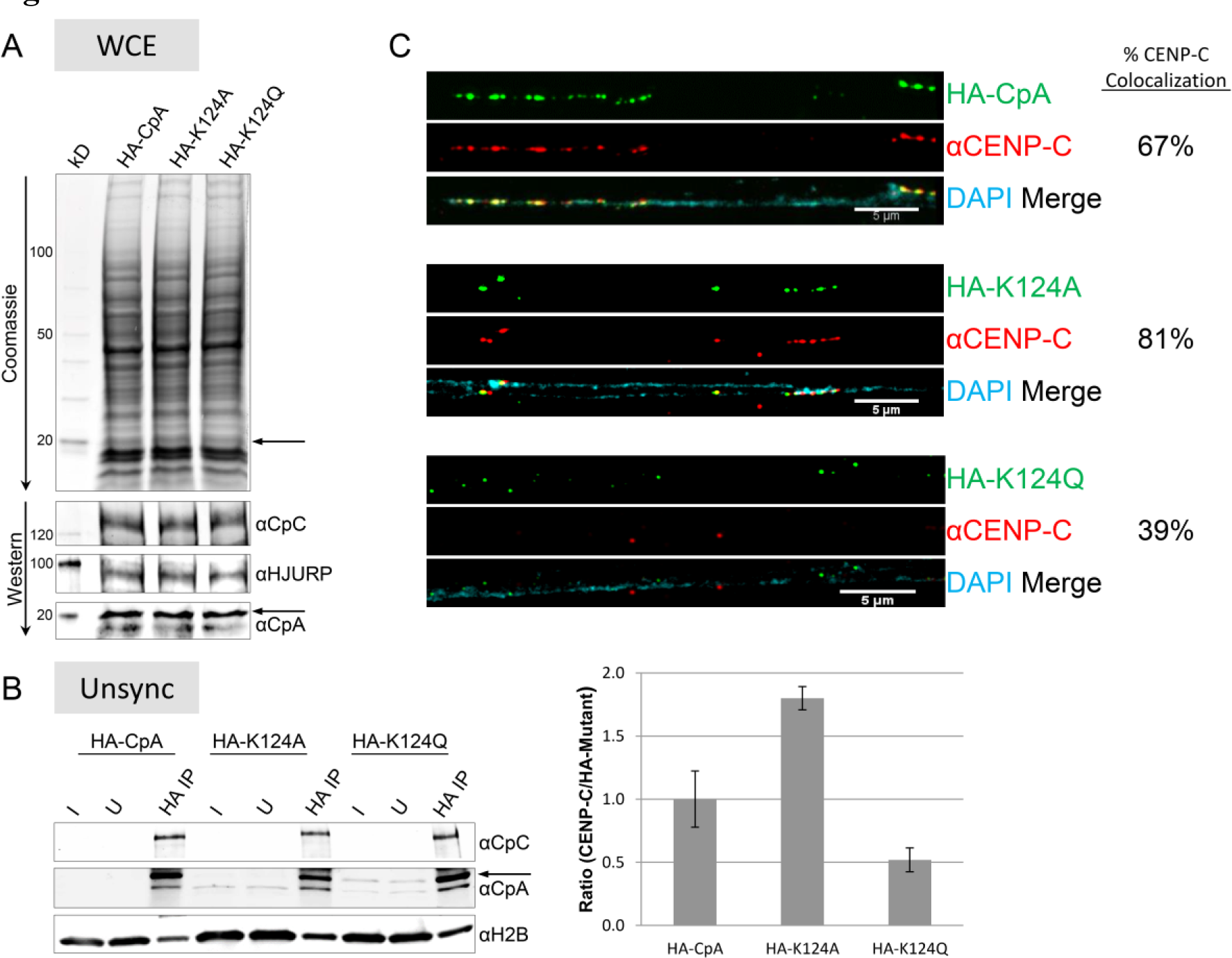
K124A/Q have altered affinity for CENP-C. **A)** Histone migration diagrammed on a traditional SDS-PAGE versus a Long TAU (LT) gel. LT gels separate histones into different combinatorial charged species, such as multi-acetylated (Ac) residues. **B)** Anti-HA ChIP was performed on medium sized chromatin arrays (see **Fig S4B**) and the amount of co-IP’ed CENP-C (CpC) was determined. Arrow indicates location of HA-tagged CENP-A/K124A/K124Q versus native CENP-A (nCENP-A) below. Three independent replicates conducted using the HA-tagged CENP-A proteins were analyzed by measuring the ratio of CENP-C/ChIP’ed HA-tagged CENP-As (wild-type or K124A/Q proteins), using LiCor Odyssey linear quantification software. Black arrow indicates HA-tagged proteins and bars indicate standard error of the mean ratio. I= Input and U=Unbound. C) Cells expressing HA-tagged CpA, K124A or K124Q mutants had their chromatin fiber extracted, and immuno-stained against HA (green) and CENP-C (red) to look for enrichment or depletion of CENP-C on the chromatin fiber (DAPI).

HA-tagged CENP-As were then chromatin immuno-recipitated (ChIP’ed), and immunoblots were performed to detect the presence and levels of CENP-C relative to levels of CENP-A/K124A/K124Q, nCENP-A normalized over histone H2B (**Fig 5B** and **Fig S4B**). We determined the ratio of native CENP-C normalized against HA-tagged K124A/Q or CENP-A in each lane. Data from three independent experiments indicate that almost twice as much unacetylatable K124A binds CENP-C compared to CENP-A (**Fig 5B**). In contrast, the binding of acetyl-mimic K124Q to CENP-C is reduced by 50% relative to CENP-A (**Fig 5B**). These data suggest K124 acetylation subtly modulates the association of CENP-C with the CENP-A chromatin fiber.

To verify this result using an independent approach, and to localize the distribution of CENP-C specifically on mutant-containing chromatin arrays, we turned to co-immunofluorescence (IF) on chromatin fibers extracted from cells expressing either HA-wildtype, HA-K124A, or HA-K124Q CENP-A. Next, we used anti-HA and anti-CENP-C antibodies to examine the association of CENP-C on HA-CENP-A positive chromatin fibers (**Fig 5C**). We scored fibers as “co-localization positive” only if >50% spots on a given contiguous single DAPI-stained chromatin fiber showed colocalization of the HA-tagged mutant with CENP-C foci; conversely, if <50% of spots on a fiber did not co-localize, such fibers were scored as “co-localization negative”. Using this metric, we observed that 12/18 (67%) wildtype HA-CpA fibers demonstrated robust colocalization with CENP-C. In contrast, 21/26 (81%) fibers displayed colocalization in the HA-K124A mutant, but only 11/28 (39%) fibers displayed robust co-localization between HA-K124Q and CENP-C. These independent results support the biochemical co-IP results above (**Fig. 5B**), namely that relative to wildtype CENP-A, CENP-A K124A has enhanced CENP-C associations, whereas CENP-A K124Q has reduced associations with CENP-C.

Curiously, we also observed K124Q fibers on which a small fraction (<10%) of K124Q tends to form discrete “clumped” structures along the chromatin fiber, while HA-CpA and HA-K124A generally form arrays wherein the foci are aligned in sequential order. Interestingly, this sub-population of CENP-A K124Q which forms large foci, appears to associate with CENP-C (**Fig. 5C**). We interpret these data to mean that if CENP-A K124Q manages to fold into superfolded beads, these structures can retain CENP-C, potentially by direct or indirect associations; whereas the larger fraction of individual spots of CENP-A K124Q dotted along the same chromatin fiber, appear not to retain CENP-C.

### Constitutive gain or loss of K124 acetylation results in modest increase in rates of mitotic errors

In previous experiments, CENP-C over-expressed in chicken DT40 cells led to higher rates of chromosomal segregation errors (Fukagawa *et al*., 1999). Since our data pointed to alterations in binding of CENP-C to mutant CENP-A, we next examined whether the presence of constitutive CENP-A K124A or K124Q correlated with any mitotic defects.

After transfection of GFP-CENP-A or GFP-K124A/Q, followed by immuno-staining with alpha tubulin, we then scored cells at anaphase to assess the frequency of multi-polar spindles and lagging chromosomes. Consistent with previous data, GFP-tagged CENP-A (CpA) cells had 14% of cells with lagging chromosomes (white arrow) (**Fig 6**). In contrast, GFP-K124A cells exhibited twice as many cells with defects, with 24% lagging, 4% multi-polar and 3% lagging + multi-polar (**Fig 6**). Similarly to the K124A mutant, K124Q cells also exhibited greater number of cells with mitotic defects, with 13% lagging, 6% multi-polar and 5% lagging + multi-polar (**Fig 6**). Although we cannot exclude that mitotic errors seen here arise from an accumulation of downstream defects, our data support previous observations that changes in CENP-C levels on CENP-A chromatin correlate with an increase in the rate of mitotic errors (Fukagawa *et al*., 1999).

**Figure 6.**
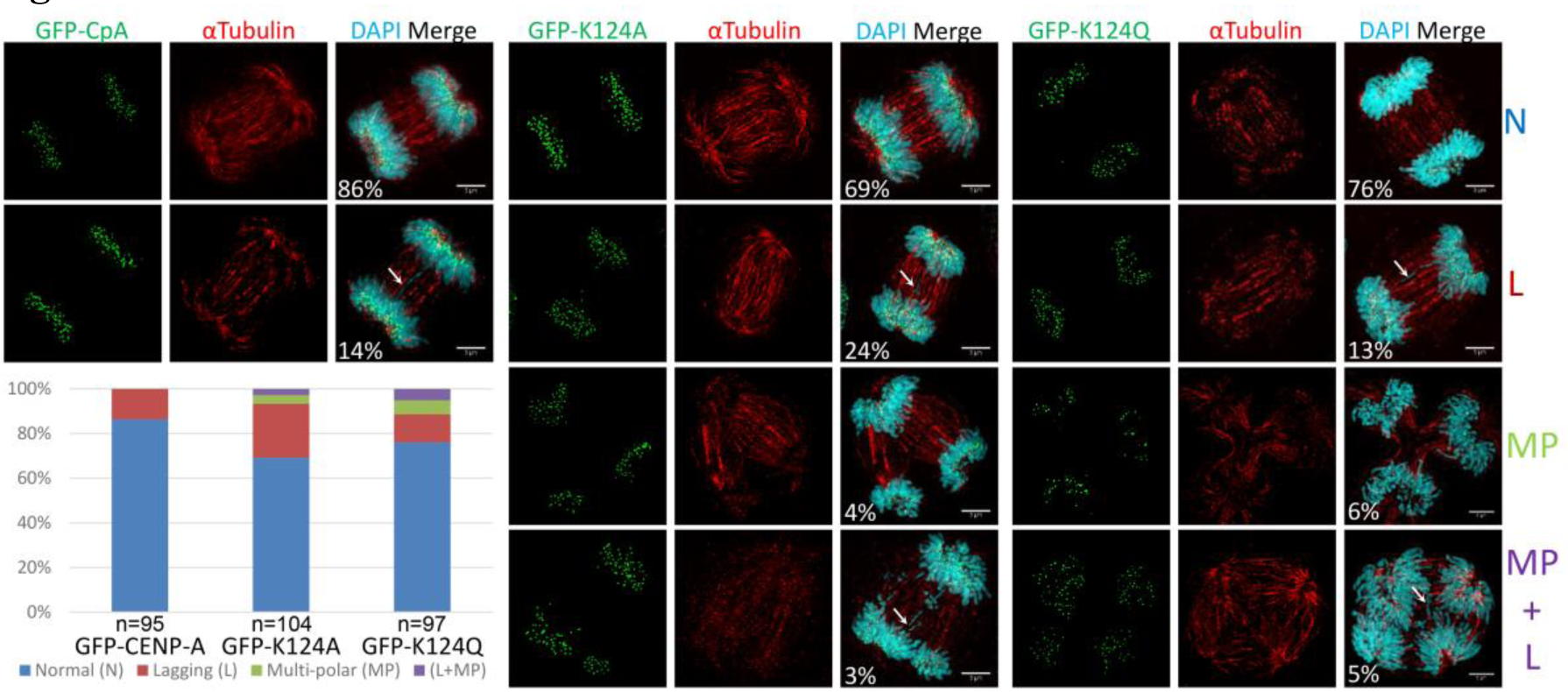
K124A/Q mutant cells have higher rates of mitotic defects compared to wild-type CENP-A. Cells were transfected with GFP-tagged CpA, K124A or K124Q, and synchronized with a double thymidine block to enrich for mitotic cells. Syncrhonized cells were immuno-stained for αTubulin, and images captured, binned and quantified into four categories: normal mitosis, mitosis with lagging chromosomes (white arrow), multi-polar spindles and multi-polar spindles + lagging chromosomes (white arrow). Percentage of cells in each category for each mutant is shown graphically and values stated in the merge panel.

### K124A/Q have altered centromeric replication timing

Native CENP-A K124ac was first discovered during G1/S, a critical junction before the onset of replication (Bui *et al*., 2012), leading us to originally suggest that K124ac might, in normal cells, contribute to centromere priming for replication (Bui *et al*., 2012; Bui *et al*., 2013). Indeed, prior work has demonstrated that a single modification within histone H3 sets the stage for accurate origin firing during replication (Prasanth *et al*., 2004; Giri *et al*., 2015). Furthermore, the computational predictions from the above experiments (**Fig 1B**) suggested that K124ac CENP-A nucleosomes had a higher potential for DNA unwrapping and sliding, which could promote CENP-A nucleosomal mobility. This in turn, could provide the mechanistic basis for our previous finding that the centromeric chromatin fiber undergoes significant decompaction at the G1/S→S transition (Bui *et al*., 2012), a result later reported in a separate study (Yu *et al*., 2015). To test whether long term gain or loss of K124ac impacted centromeric replication, we returned our attention to S phase.

We synchronized cells, released them at S phase, and staged cells to well-established early, mid and late S-phase time points (Dimitrova and Berezney, 2002) (diagramed in **Fig S2B**). We used EdU staining to detect replication foci (Nowakowski *et al*., 1989), coupled to CENP-B staining, since CENP-B is a well established marker for independent and sequence-specific binding to human centromeric DNA. To quantify replication timing, we scored the number of EdU+/CENP-B+ foci over the total number of CENP-B foci in K124A, K124Q, or wildtype CENP-A expressing cells. We observed 25-30% EdU+/CENP-B+ spots in wild-type CENP-A in early and mid S phases, with more of the EdU+/CENP-B+ colocalizing in late S-phase (37%) (**Fig 7**). These data are consistent with previous documentation of late replication of centromeres (Ten Hagen *et al*., 1990), with a minor fraction of centromere replication distributed throughout S-phase (Weidtkamp-Peters *et al*., 2006; Erliandri *et al*., 2014). In contrast, K124A mutants had only 15% EdU+/CENP-B+ co-stained foci in early and mid S phase. By late S phase, replicating centromeres in the K124A mutant cells were restored to wild-type levels of 37%. We performed FACS analysis of the mutants to examine whether any genome-wide ploidy changes could be detected, finding that cells progress normally through the cell cycle with no ploidy defects in K124A or K124Q cells (**Fig S4A**). These data suggest that the K124A mutant has a modest replication defect, in which it is slow to initiate centromeric replication in early S, but which eventually restores to wild type levels before the next mitosis.

**Figure 7.**
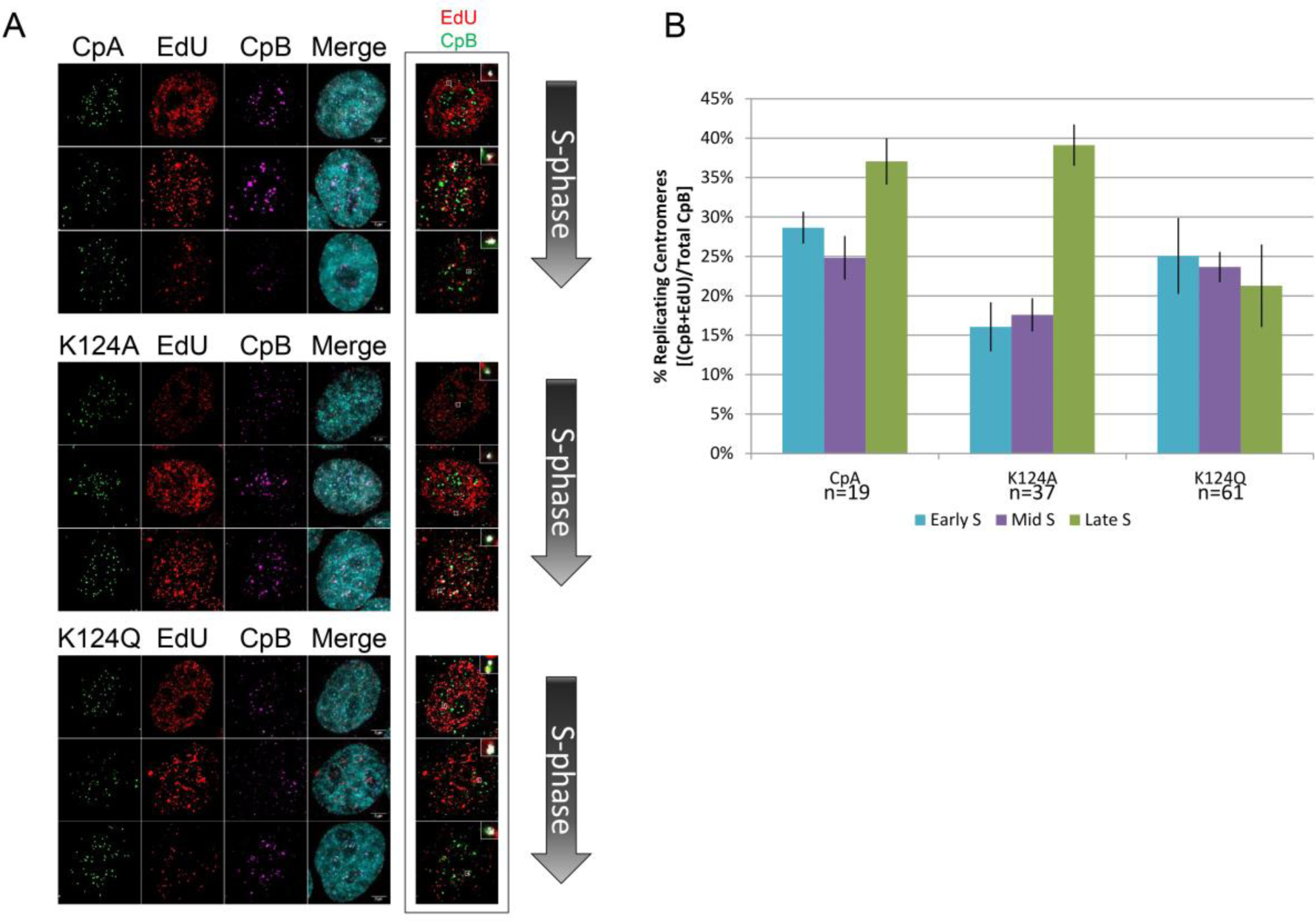
CENP-A mutations in K124 display altered replication timing of centromeric foci. **A)** EdU pulsed cells containing wild-type CENP-A (CpA) and K124A/Q were co-stained with CENP-B (CpB) to assess percentage of centromeric replicating origins at early, mid and late S-phases. 5 um scale bars are located in the bottom right of the merge panel. Boxed to the right is an example of automated co-localization analysis using ImageJ which was used to determine the fraction of co-localizing CpB and EdU foci (in white) with EdU+/CpB+ insets to show co-localization. **B)** A graphical representation of the percentage of centromeric origins co-stained with EdU (CpB+EdU) over the total number of centromeric (marked by CpB) foci.

Conversely, we observed that K124Q displayed the opposite type of centromeric replication defect. In contrast to wildtype CENP-A cells, in which majority of CENP-B foci co-localized with EdU mostly at late S phase, in K124Q cells, EdU+/CENP-B co-localizing foci shift toward early→mid-S replication, with EdU+/CENP-B+ foci distributed equally over early, mid and late S-phase (**Fig 7B**). Thus, constitutive gain- or loss- of K124 acetylation status correlates with alterations in the distribution of centromeric replication timing.

### CENP-A K124 switches from acetylation at G1/S to monomethylation at S-phase

The replication defects noted in K124Q above, wherein centromeric foci are modestly advanced in replication timing, would suggest that K124ac must presumably be removed by early S phase, in order for centromeric replication to progress correctly. To address whether acetylation at K124ac is removed, or replaced by another modification at S phase, we utilized a native CENP-A purification strategy coupled to MS/MS approach as in previous studies (Waterborg *et al*., 1995; Hake *et al*., 2006; Shechter *et al*., 2007; Bui *et al*., 2012), adding a super-resolving, double long Triton-Acid-Urea (DLτ or DL-TAU) gel (see Methods, and Nuccio, Bui et al Methods in Enzymology Proteomics B, 2017). Unlike traditional long TAU gels where all histones remain in the gel, in the case of the DLτ, we run the majority of core histone proteins off the gel, leaving behind only histone H2A at the bottom as marker, resulting in CENP-A species distributed throughout the middle of the gel (**Fig 8A** and **Fig S5**). This permits enhanced enrichment of CENP-A species while reducing promiscuous core histone contaminants during excision of gel slices for MS/MS analysis. This strategy also helps to assure that any antibody-based staining used to detect modifications reflects true CENP-A containing species rather than mis-interactions with H3. The use of TAU-gels is especially important when studying CENP-A modifications, because many commercially available CENP-A antibodies display frequent background interactions with H3 (*unpublished data*), exacerbated by the fact that H3 is present in hundred-fold molar excess over CENP-A in chromatin extracts, is heavily modified itself, and co-migrates with CENP-A on standard SDS-PAGE gels.

**Figure 8.**
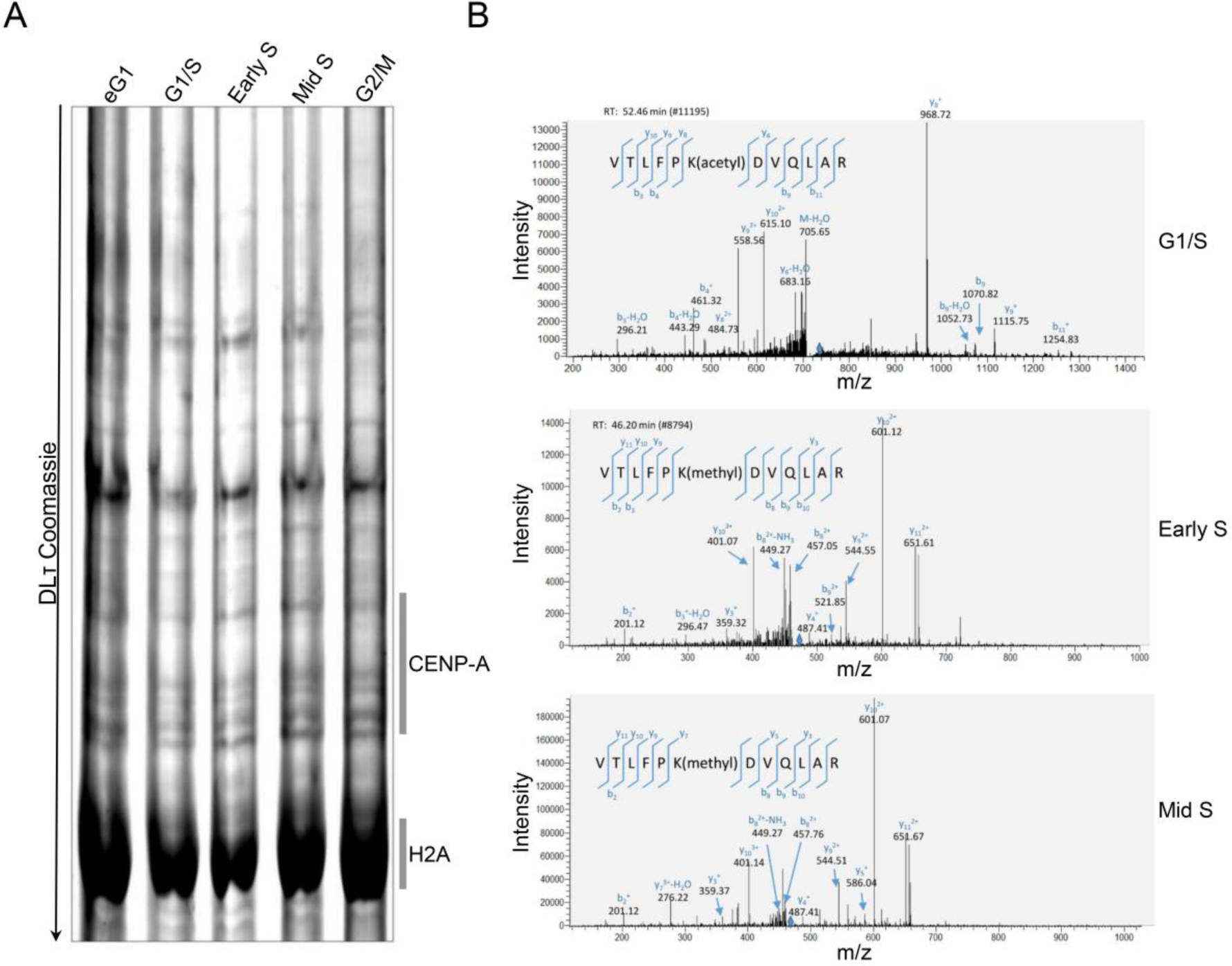
Endogenous CENP-A K124 is acetylated at G1/S but monomethylated at early and mid S-phases. **A)** G1/S, eS and mid-S Chromatin bound histones were isolated, resolved on a Double Long TAU (DLτ) gel, stained with Coomassie, and endogenous CENP-A species excised for subsequent analysis by mass spectrometry. **B)** A peptide containing acetylated lysine 124 was observed in the G1/S. The representative MS/MS spectrum showing CENP-A K124 acetylated in the peptide “VTLFPK(acetyl)DVQLAR” is shown on the bottom left. Location of the parent peptide ion (m/z = 714.90, charge = +2) prior to fragmentation is indicated in each spectrum with a blue diamond. Peptide fragmentation ions identified are indicated in the spectra and on the peptide sequence. The masses of ions b9, b11, y8, y9, and y10 are diagnostic of K124 acetylation. The peptide containing mono-methylated lysine 124 was observed in the early S and mid-S phase – the representative MS/MS spectrum showing CENP-A K124 methylated in the peptide “VTLFPK(methyl)DVQLAR” is shown on the middle and top left. Location of the parent peptide ion (m/z = 466.60, charge = +3) prior to fragmentation is indicated in each spectrum with a blue diamond. Peptide fragmentation ions identified are indicated in the spectra and on the peptide sequence. The masses of ions b8, b10, y7, y9 and y10 are diagnostic of K124 methylation.

Mass spectra of CENP-A containing bands from these DLτ gels confirm our earlier findings that CENP-A is acetylated at K124 during G1/S (Bui *et al*., 2012). Interestingly, this modification is not detected in either early or mid S-phases (**Fig 8B**). Instead, in both early and mid S phases, we detected robust methylation of K124 (**Fig 8B**, **Tables 1**–**3**). Conversely, K124me was not detected in the G1/S CENP-A bands. These data suggest a cyclical nature to modifications of K124, so that permanent gain or loss of acetylation at K124 likely disrupts the cycling of K124ac → K124me, which, in turn, impacts centromere dynamics during replication and subsequent mitoses.

**table 1.** Mass spec fragment ion table for G1/S phase

| #1 | b<sup>+</sup> -<br>Theoretical | b<sup>+</sup> -<br>Experimental | b<sup>2+</sup> -<br>Theoretical | b<sup>2+</sup> -<br>Experimental | Seq. | y<sup>+</sup> -<br>Theoretical | y<sup>+</sup> -<br>Experimental | y<sup>2+</sup> -<br>Theoretical | y<sup>2+</sup> -<br>Experimental | #2 |
| --- | --- | --- | --- | --- | --- | --- | --- | --- | --- | --- |
| 1 | 100.0757 |  | 50.54149 |  | V |  |  |  |  | 12 |
| 2 | 201.12338 |  | 101.06533 |  | T |  |  | 665.37993 | 665.66638 | 11 |
| 3 | 314.20745 | 314.2298 | 157.60736 |  | L | 1228.7049 | 1228.83008 | 614.85609 | 615.0849 | 10 |
| 4 | 461.27587 | 461.30969 | 231.14157 |  | F | 1115.62083 | 1115.7052 | 558.31405 | 558.50128 | 9 |
| 5 | 558.32864 | 558.50128 | 279.66796 |  | P | 968.55241 | 968.66461 | 484.77984 | 485.78 | 8 |
| 6 | 728.43417 |  | 364.72072 |  | K-Acetyl | 871.49964 | 871.6875 | 436.25346 |  | 7 |
| 7 | 843.46112 | 843.41919 | 422.2342 |  | D | 701.3941 | 701.29291 | 351.20069 |  | 6 |
| 8 | 942.52954 | 942.66 | 471.76841 | 471.3 | V | 586.36715 | 586.47998 | 293.68721 |  | 5 |
| 9 | 1070.58812 | 1070.74 | 535.7977 |  | Q | 487.29873 | 487.38043 | 244.153 |  | 4 |
| 10 | 1183.67219 | 1183.84912 | 592.33973 | 592.63 | L | 359.24015 | 359.27 | 180.12371 |  | 3 |
| 11 | 1254.70931 | 1254.79114 | 627.85829 | 627.32 | A | 246.15608 | 246.14738 | 123.58168 |  | 2 |
| 12 |  |  |  |  | R |  |  | 88.06312 |  | 1 |

**Table 2.** Mass spec fragment ion table for early S phase

| #1 | b<sup>+</sup> -<br>Theoretical | b<sup>+</sup> -<br>Experimental | b<sup>2+</sup> -<br>Theoretical | b<sup>2+</sup> -<br>Experimental | b<sup>3+</sup> -<br>Theoretical | b<sup>3+</sup> -<br>Experimental | Seq. | y<sup>+</sup> -<br>Theoretical | y<sup>+</sup> -<br>Experimental | y<sup>2+</sup> -<br>Theoretical | y<sup>2+</sup> -<br>Experimental | y<sup>3+</sup> -<br>Theoretical | y<sup>3+</sup> -<br>Experimental | #2 |
| --- | --- | --- | --- | --- | --- | --- | --- | --- | --- | --- | --- | --- | --- | --- |
| 1 | 100.08 |  | 50.54 |  | 34.03 |  | V |  |  |  |  |  |  | 12 |
| 2 | 201.12 | 201.13 | 101.07 |  | 67.71 |  | T | 1301.75766 |  | 651.38247 | 651.58466 | 434.59074 | 434.88 | 11 |
| 3 | 314.21 | 314.26 | 157.61 |  | 105.41 |  | L | 1200.70998 |  | 600.85863 | 601.03448 | 400.90818 | 401.00195 | 10 |
| 4 | 461.28 | 460.65 | 231.14 |  | 154.43 |  | F | 1087.62591 |  | 544.31659 | 544.31 | 363.21349 | 363.26254 | 9 |
| 5 | 558.33 | 558.50 | 279.67 | 279.13 | 186.78 |  | P | 940.55749 |  | 470.78238 |  | 314.19068 | 314.26239 | 8 |
| 6 | 700.44 |  | 350.72 | 350.18 | 234.15 |  | K-Methyl | 843.50472 |  | 422.256 | 422.16428 | 281.83976 | 281.40979 | 7 |
| 7 | 815.47 |  | 408.24 | 408.10 | 272.49 | 272.21 | D | 701.3941 |  | 351.20069 | 351.17957 | 234.46955 |  | 6 |
| 8 | 914.53 |  | 457.77 | 458.04 | 305.52 |  | V | 586.36715 | 585.92181 | 293.68721 | 294.14 | 196.12723 |  | 5 |
| 9 | 1042.59 |  | 521.80 | 521.63 | 348.20 |  | Q | 487.29873 | 487.35 | 244.153 | 244.11246 | 163.10443 |  | 4 |
| 10 | 1155.68 |  | 578.34 | 578.60 | 385.90 | 385.20 | L | 359.24015 | 359.29077 | 180.12371 |  | 120.41823 |  | 3 |
| 11 | 1226.71 |  | 613.86 | 613.82 | 409.58 | 409.25 | A | 246.15608 | 246.22052 | 123.58168 |  | 82.72354 |  | 2 |
| 12 |  |  |  |  |  |  | R | 175.11896 | 175.16 | 88.06312 |  | 59.0445 |  | 1 |

**Table 3.** Mass spec fragment ion table for mid S phase

| #1 | b<sup>+</sup> -<br>Theoretical | b<sup>+</sup> -<br>Experimental | b<sup>2+</sup> -<br>Theoretical | b<sup>2+</sup> -<br>Experimental | b<sup>3+</sup> - Theoretical | b<sup>3+</sup> -<br>Experimental | Seq. | y<sup>+</sup> - Theoretical | y<sup>+</sup> -<br>Experimental | y<sup>2+</sup> -<br>Theoretical | y<sup>2+</sup> -<br>Experimental | y<sup>3+</sup> -<br>Theoretical | y<sup>3+</sup> -<br>Experimental |
| --- | --- | --- | --- | --- | --- | --- | --- | --- | --- | --- | --- | --- | --- |
| 1 | 100.08 |  | 50.54 |  | 34.03 |  | V |  |  |  |  |  |  |
| 2 | 201.12 | 201.16 | 101.07 |  | 67.71 |  | T | 1301.76 |  | 651.38 | 651.67 | 434.59 | 434.74 |
| 3 | 314.21 |  | 157.61 |  | 105.41 |  | L | 1200.71 |  | 600.86 | 601.07 | 400.91 | 401.14 |
| 4 | 461.28 | 460.52 | 231.14 |  | 154.43 |  | F | 1087.63 |  | 544.32 | 544.51 | 363.21 |  |
| 5 | 558.33 |  | 279.67 |  | 186.78 |  | P | 940.56 |  | 470.78 |  | 314.19 |  |
| 6 | 700.44 |  | 350.72 |  | 234.15 |  | K-Methyl | 843.50 |  | 422.26 |  | 281.84 |  |
| 7 | 815.47 |  | 408.24 |  | 272.49 |  | D | 701.39 |  | 351.20 |  | 234.47 |  |
| 8 | 914.53 |  | 457.77 | 457.76 | 305.52 |  | V | 586.37 | 586.04 | 293.69 |  | 196.13 |  |
| 9 | 1042.59 |  | 521.80 | 521.75 | 348.20 |  | Q | 487.30 | 487.34 | 244.15 |  | 163.10 |  |
| 10 | 1155.68 |  | 578.34 |  | 385.90 |  | L | 359.24 | 359.37 | 180.12 |  | 120.42 |  |
| 11 | 1226.71 |  | 613.86 |  | 409.58 |  | A | 246.16 |  | 123.58 |  | 82.72 |  |
| 12 |  |  |  |  |  |  | R | 175.12 |  | 88.06 |  | 59.04 |  |

### HATs and HDACs influencing the acetylation status of native CENP-A K124

To investigate HATs and HDACs that might influence modification of K124 in native CENP-A, our strategy again turned to Lτ gels that are capable of separating histone proteins based on charge and hydrophobicity. This gel chemistry was developed over two decades ago in the chromatin field, and is widely recognized as a robust method to exclusively study histone variants and their post-translational modifications (Waterborg *et al*., 1995; Hake *et al*., 2006; Shechter *et al*., 2007). As histone residues become postranslationally modified, such as the prescence of a lysine acetylation, the positive to neutral charge of the residue induces a positive shift on the Lτ gel. As the schematic in **Fig 9A** demonstrates, unlike traditional SDS-PAGE gels, an Lτ gel is sufficient to separate core histones into different species. This further separation of the histones into different modified subspecies increases confidence in our analysis as antibodies are known to have cross-reactivity. For examples, multiple species of histone H4 were previously identified using this kind of TAU gel (Earley *et al*., 2007), demonstrating that this gel chemistry is optimized to examine histone modifications. We developed a custom rabbit polyclonal antibody against CENP-A K124ac. Using recombinant histone proteins including CENP-A, H4, synthetic CENP-A K124ac, H4K79ac, H2A and H2B (gifts from Dr. Jennifer Ottesen), we observed a high affinity for the polyclonal antibody against the recombinant CENP-A K 124ac protein (**Fig 9A**) specifically on TAU-WB blots.

**Figure 9.**
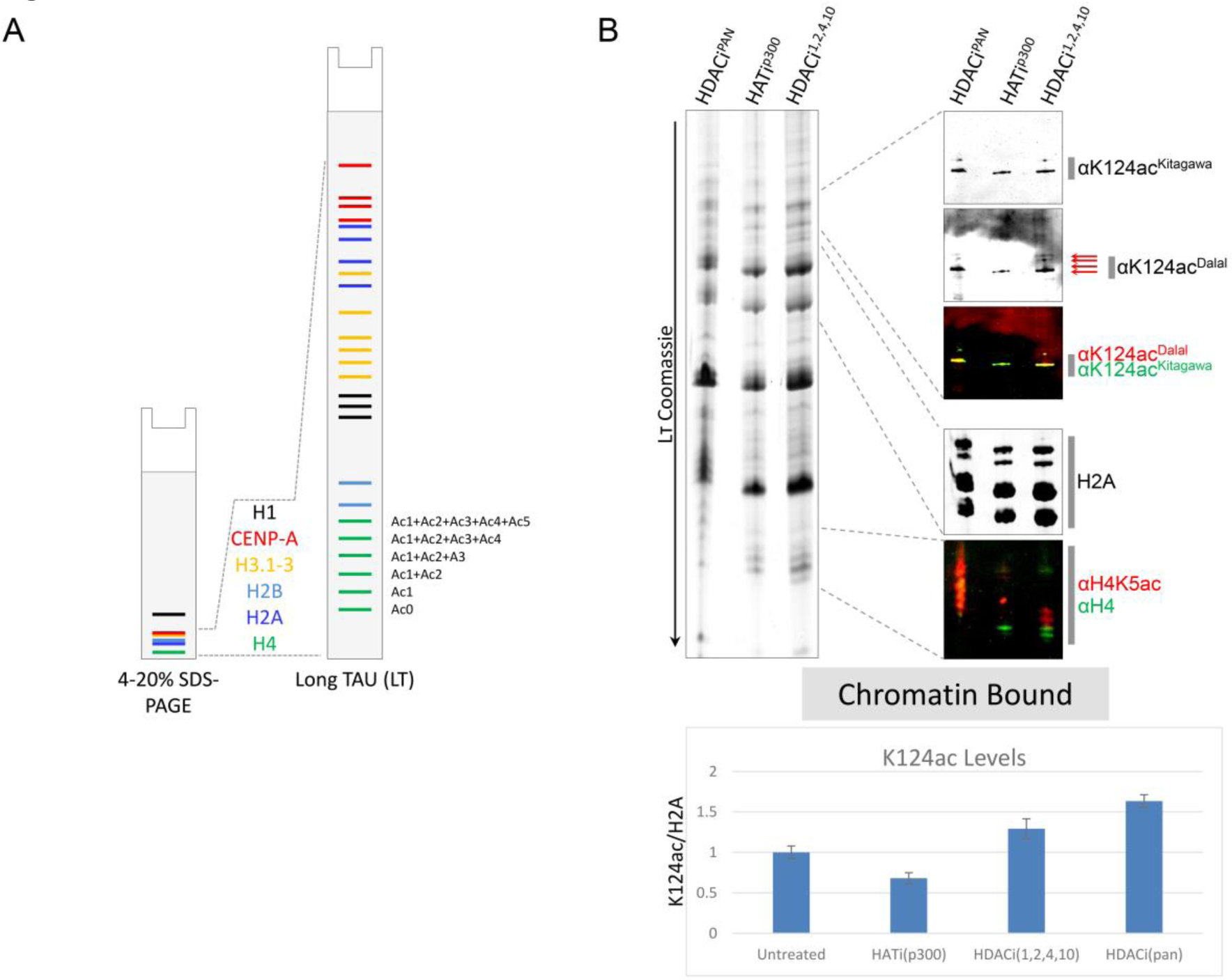
Histone acetyltransferase p300 targets CENP-A K124 for acetylation. **A)** Cells were treated for 20 hrs with C646 (HAT inhibitor against p300 = HATi^p300^) or trichostatin A/TSA (pan HDAC inhibitor = HDACi^PAN^), and whole cells extracted and ran on an SDS-PAGE or L-TAU gel, followed by Western analysis. Recombinant unmodified CENP-A and CENP-A K124ac proteins were also loaded to determine antibody specificity, and levels of K124ac were normalized against total proteins and relative to untreated = 1. **B)** Chromatin bound histones were purified from nuclei and ran on an L-TAU gel followed by Western analysis. In addition to treating cells with C646 or TSA, cells were also treated with quisinostat, a weak pan HDAC inhibitor that targets HDAC 1, 2, 4 and 10 (HDACi^1,2,4,10^). Average K124ac levels were normalized against total histone H2A and plotted relative to untreated = 1.

To examine the role of various HAT/HDAC complex on CENP-A K124ac, we treated cells with well-characterized histone acetyltransferase (HAT) and histone deacetylase (HDAC) inhibitors. C646 is a HAT inhibitor that targets p300 (HATi^p300^), trichostatin A (TSA) is a potent pan HDAC inhibitor (HDACi^PAN^), and quisinostat 2HCl is a weak pan inhibitor that targets predominantly HDAC1, 2, 4 and 10 (HDACi^1,2,4,10^). P300 was predicted as a strong HAT target because of a previous study, in which co-localization of p300 with CENP-A was observed at centromeres (Shimura *et al*., 2011). Furthermore, it has been shown that tethering of a p300 catylytic domain induces assembly of CENP-A onto alphoid DNA (Ohzeki *et al*., 2012; Shono *et al*., 2015), in which it was suggested that acetylation of CENP-A was required for its stable retention on chromatin. Cells were treated with control DMSO, or DMSO with 20 μM C646 or 200 nM TSA, for 20 hrs, harvested, and subsequent purification of either whole cell extracts (WCE), or chromatin-bound histones using hydroxyapatite-elution (Bui *et al*., 2012). In both cases, WCE, or chromatin-bound, histones were probed on an SDS-PAGE Western for GAPDH and also on an L-TAU Western using the CENP-A K124ac antibody, CENP-A, H2A and H4. Because it has been previously shown that inhibition of p300 by C646 greatly reduces the level of H4K5ac (Henry *et al*., 2015), we used H4K5ac as a control to confirm the drug functions correctly in our hands. As expected, H4K5ac levels diminish significantly in the p300 inhibitor (C646) treatment compared to untreated, suggesting effective inhibition of p300 (**Fig S5B**). Probing with the CENP-A K124ac antibody reveals a higher affinity for the recombinant CENP-A K124ac protein compared to unmodified recombinant CENP-A (**Fig S5B**). However, in WCE, after normalizing CENP-A K124ac with total histones on the TAU gell, we observed that CENP-A K124ac levels were only modestly reduced (**Fig S5B**). We next examined chromatin bound histones from nuclei, using canonical histone H2A as a loading and normalizing control over untreated. As expected, in the p300 inhibition lane, control H4K5ac levels were significantly reduced. We next co-stained our TAU-WB blots with our custom K124ac antibody, along with a second antibody that also recognizes CENP-A K124ac (a gift from Dr. Katsumi Kitagawa). Reassuringly, both K124ac antibodies recognized the same CENP-A species on TAU-WB (**Fig 9B**). Unlike in WCE, chromatin bound levels of CENP-A K124ac in the HAT and HDAC inhibited lanes were markedly different than the untreated sample. CENP-A K124ac levels were reduced by about 40% (compared to untreated) in the p300 inhibited cells, while there was a 30% and 60% increase from the quisinostat and pan HDAC TSA treatment, respectively (**Fig 9B**). Thus, our data suggest that p300 likely functions in the K124acetylation pathway. Our evidence also supports the idea that multiple HDACs may be involved in removing the acetylation of K124, because we observed reduction of K124ac species in the pan and HDAC-specific inhibitor lanes (**Fig 9B**). Taken together, these data suggest p300 functions on chromatin bound CENP-A to promote its acetylation.

## DISCUSSION

Despite decades of intensive biochemical studies, how modifications impact the histone core and subsequent function have been difficult to dissect. In recent years, elegant *in vivo* and *in vitro* experiments have shown that these core modifications can exert profound effects on nucleosome behavior and subsequent chromatin changes (Schneider *et al*., 2006; Manohar *et al*., 2009; Shimko *et al*., 2011; Tropberger *et al*., 2013; Brehove *et al*., 2015). In previous work, we reported that during G1/S, human centromeric chromatin becomes more accessible, and that CENP-A nucleosomes occupy a transitionary state concurrent with internal histone fold domain acetylations at CENP-A K124 and H4K79 (Bui *et al*., 2012). We originally suggested that CENP-A K124ac and H4K79ac were responsible for a structural change, altering from stable tetrameric to stable octameric nucleosomes inferred by AFM analysis. However, an alternative interpretation of changes in CENP-A dimensions that is supported by current computational and experimental data, is that the CENP-A octamer, in the absence of modifications or binding partners, is dynamic and flexible, capable of undergoing shearing motions which may expose the internal core of the protein octamer (Hasson *et al*., 2013; Falk *et al*., 2015; Winogradoff *et al*., 2015b).

In this work, we probed the effects of histone core modifications of CENP-A using a combinatorial approach. Computational modeling demonstrates that CENP-A K124ac and H4K79ac collaborate to loosen the DNA at four symmetric contact points within the nucleosome, which is predicted to promote access to the nucleosomal DNA whilst simultaneously rigidifying the protein core. *In silico* results also suggest that dual acetylation of CENP-A K124 and H4K79 serves an un-anticipated role in restraining the C-terminus of CENP-A so that it is less accessible than in the unmodified CENP-A NCP (**Fig 1**–**3**). One outcome of this “locking” of the CENP-A C terminus is a reduced binding to CENP-C *in silico*.

In subsequent biological experiments, by expressing mimics of acetylated K124 (K124Q) or unacetylatable K124 (K124A) CENP-A, we observed that both these mutants have altered CENP-C binding (**Fig 5**). In the unacetylatable mimic (K124A), CENP-C was strongly bound, whereas in the acetylated mimic, CENP-C binding dropped to 50% of wildtype. These results are supported by chromatin fiber co-IF, which likewise shows a reduced association of CENP-C on K124Q fibers, and a relatively increased association on K124A fibers. Consistent with direct or downstream accumulated changes in CENP-C binding, K124A and K124Q cells display a relative increase in mitotic defects relative to wild-type CENP-A cells. These data are consistent with earlier experiments documenting that titration of CENP-C levels lead to higher mitotic error rates (Fukagawa *et al*., 1999).

Our experiments investigating the role of K124ac at replication shows that in cells expressing the acetyl mimic, K124Q, centromeric foci are relatively ahead in replication timing, whereas in cells expressing the unacetylatable K124A, centromeric replication is delayed (**Fig 7**). The mechanistic basis for this observation remains under investigation. The simplest possibility is that the “opening” of the CENP-A chromatin fiber is prerequisite to replication. Not mutually exclusive to this hypothesis, it is feasible that defects in ORC binding contribute to the delay in replication timing. The pre-Replication Complex (pre-RC) consists ORC1 through ORC6 (DePamphilis, 2003), of which, only ORC2 has been shown to specifically associate with centromeres at S phase (Prasanth *et al* 2004). Precedence comes from recent work which has demonstrates that centromeric replication timing is exquisitely sensitive to the state of the chromatin fiber: depletion of the key inner kinetochore protein CENP-B alters the chromatin landscape at centromeres, and enhances the binding of ORC2 to centromeric origins (Erliandri *et al*., 2014). We speculate that the centromeric replication defects observed in our study above (**Fig 7**) is similarly underpinned molecularly by a combination of chromatin fiber accessibility, and mis-timing of the interaction between K124A/Q mutant proteins at the centromeric fiber with potentially one or more of the pre-RC components. Importantly, the defects in replication timing of the permanently acetylated state K124Q, also point to the importance of *removing* the acetylation by early to mid S-phase. Here, our experimental MS/MS data demonstrate that CENP-A K124 acetylation is replaced by a robust CENP-A K124 methylation at S phase (**Fig 8**).

Precedence for histone core domain methylation in regulating replication timing exists. Methylation of K79 in canonical H3 is critical, and limits genome-wide DNA replication to once per cell cycle, thereby preventing over-replication (Fu *et al*., 2013). In our mutants, neither CENP-A K124A and K124Q can undergo methylation, breaking the potential cycle of G1/S phase K124ac→S phase K124me (**Fig 10A-B**). We hypothesize that one putative function of K124me may be to stabilize the nucleosomal DNA, either by stabilizing re-binding of CENP-C, or, not mutually exclusive, potentially to inhibit rebinding of preRC proteins to the newly replicated CENP-A chromatin, thereby presenting a barrier to over-replication (**Fig 10C**). A strong possibility based on the known function of H3K122ac in facilitating euchromatic transcription informs this hypothesis further. We speculate that CENP-A K124Me increases CENP-C binding, closing the chromatin fiber, which could inhibit centromeric transcription in G2. The potential loss of transcription-mediated eviction of CENP-A in the absence of new CENP-A loading, would be deleterious, since the only other protein present to load chromatin in G2 is the transcription-coupled histone H3.3. In this scenario, removing acetylation from CENP-A would also be important for downstream events in G2/M. Thus, subsequent mitotic defects, which may be linked either directly or indirectly to CENP-C distribution in these mutant states of CENP-A (**Fig 5**), might also reflect the importance of binding of CENP-C to CENP-A K124me nucleosomes. Consequently, identifying whether a unique species of CENP-A K124me bound CENP-C nucleosomes exist, and serve a biological function, such as modulating centromere transcription in G2, is an ongoing extension of the current work. Additional future experiments that are necessary include dissecting how HATS like p300, which, as shown above (**Fig 9**) contributes to CENP-A’s acetylation status, are recruited to the centromeric fiber, and whether it targets other candidate lysines (e.g K56, K64), and identifying the HMT responsible for methylating CENP-A K124. Excitingly, on going *in vitro* dissections using recent successful snythetically engineered CENP-A K124ac (Yu *et al*., 2016) coupled to H4K79ac proteins, will determine in vitro whether acetylated CENP-A nucleosomes encode a chromatin fiber more susceptible to sliding, and whether such a fiber has altered binding affinity to CENP-C.

**Figure 10.**
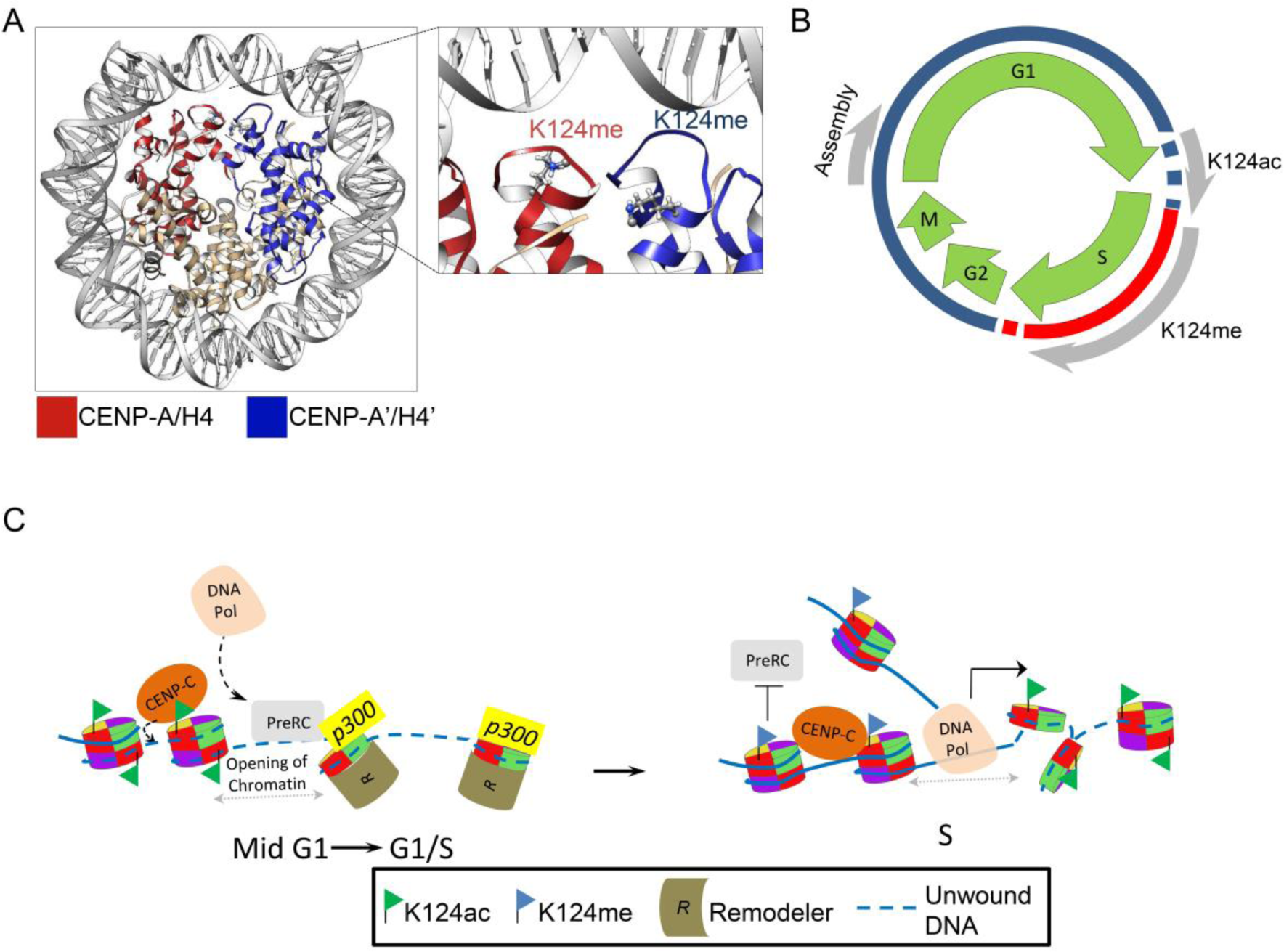
A model depicting how cyclical switching in CENP-A K124 acetylation and methylation might affect centromeric replication dynamics. **A)** Modeling of methylation on CENP-A K124. **B**) Cell cycle progression of events that occur on the CENP-A chromatin and on the protein including acetylation at G1/S and methylation at S phase. **C)** Current model encompassing the dynamics of K124 modification on the effect of replication and kinetochore protein CENP-C binding.

Taken together, in this report, we systematically dissect the role of a single modification in the CENP-A nucleosome *in silico* and *in vivo,* finding it to be involved in regulating protein CENP-C distribution on the modified CENP-A chromatin fiber, mitotic integrity, and centromere replication timing. We emphasize that delays in centromere replication timing are not permanent, as most cells seem to recover by late S phase. Indeed, majority of the mutant cells survive in the presence of K124A and K124Q, and, despite the small but measurable increase in mitotic defects, we did not observe persistent deleterious effects. Thus, while acetylation of CENP-A K124, in our view, does not play a deterministic role, it does subtly contribute to centromere dynamics at the level of replication timing, and mitotic integrity. Consequently, these data open a new avenue of investigation into how covalent modifications of histone variant nucleosomes can modulate epigenetic regulation of their cognate loci.

## MATERIALS AND METHODS

### Simulation Protocol

All-atoms molecular dynamics (MD) simulations were performed with software suite Gromacs 5.0.4 (Bekker *et al*., 1992). The force field employed to model nucleosomes was amber99SB*-ILDN (Best and Hummer, 2009; Lindorff-Larsen *et al*., 2010) for proteins, amber99SB parmbsc0 (Perez *et al*., 2007) for DNA, ions08 (Joung and Cheatham, 2008) for ions, and the TIP3P (Jorgensen *et al*., 1983) water model.

Two nucleosomal systems were built for simulation: the acetyl-lysine CENP-A nucleosome and then unmodified CENP-A nucleosome as control. First, the CENP-A nucleosome was built with PDB ID: 3AN2^1^—resolved to 3.60 Å—as the starting structure. Unresolved 3AN2 residues Thr 79 through Asp 83 of CENP-A’, Chain E, were built with MODELLER (Marti-Renom *et al*., 2000). During energy minimization of this constructed region, one residue in the n-terminus and c-terminus directions was unconstrained. Additionally, heterogen selenomethionine residues were altered to methionine through a single atom mutation from Se to S. As a control, the 146 base pair α-satellite DNA of PDB ID: 3WTP (Arimura *et al*., 2014) was aligned onto 3AN2 using the CE algorithm (Shindyalov and Bourne, 1998) of PyMOL (The PyMol Molecular Graphics System). To further study the effect of acetylation on CENP-C binding, the CENP-C fragment (Kato *et al*., 2013) was docked onto each system using the CE algorithm (Shindyalov and Bourne, 1998) and simulated for an additional microsecond retaining identical simulation protocols. This is the first known all-atom detail structural analysis of CENP-C bound to CENP-A.

From this initial structure, the Gromacs tool pdb2gmx was used to assign charges to residues at biological pH: a charge of +1 on Lys and Arg, 0 for Gln, −1 for Asp and Glu, and His with hydrogen on the epsilon nitrogen. Then, a rectangular cuboid box was created such that boundaries were a minimum distance of 1.5 nm from the unsolvated system. Next, Na+ and Cl-ions were introduced to neutralize the system charge and additionally model an ionic concentration to 150 mM. For both preproduction and production runs, periodic boundary conditions were employed. Electrostatics were handled with the Particle Mesh Ewald method and Verlet cutoff scheme. For the non-bonded interaction shift functions, Coulombic and Van der Waals potentials had a cutoff distance at 1.0 nm. Hydrogen bonds were constrained with the LINCS algorithm.

The CENP-A nucleosome system was energy minimized using steepest descent to a maximum energy of 100 kJ/mol. The systems were then equilibrated in multiple steps. First, the systems were heated to 300 K for 2000 ps. During this step, DNA was restrained with K = 1000 kJ mol^−1^ nm^−2^ in the Canonical ensemble (NVT). For the next thermal equilibration at 300K for 2000 ps, both DNA and protein had weak harmonic position restraints, KCENP-A = 2.5e-5 kJ mol^−1^ nm^−2^, to hinder global rotational motions. Lastly, pressure was equilibrated for 1500 ps in the Isothermal-isobaric, NPT, ensemble at 300 K and 1.0 bar with KCENP-A.

This system was ran for 1 μs at 300 K. Temperatures were V-rescaled with the modified Berendsen thermostat (Berendsen *et al*., 1984) with a time constant of 1.0 ps. System pressures were regulated with the Parrinello-Rahman barostat (Parinello and Rahman, 1981) at 1.0 bar and a time constant of 2.0 ps. The simulations’ time step size was 2 fs. Coordinates, velocities, and energies were saved every 2 ps. Non-bonded neighbors lists were updated every 20 fs.

After the CENP-A nucleosome was run for 1 μs, the final structure was acetylated in four histone core locations: K124 of CENP-A and CENP-A’, and K79 of H4 and H4’. The partial charges assigned to acetyl-lysine atoms were calculated quantum mechanically as described previously (Winogradoff *et al*., 2015a). The new amino acid type for acetyl-lysine, KAC, was added to amber99SB*-ILDN ^2, 3^. Both the acetyl-lysine CENP-A system and the control CENP-A system were simulated for an additional 1 μs as described above. For subsequent analysis, trajectories were truncated to remove the first 600 ns to account for additional system equilibration during production runs.

### Analysis of trajectories

After truncating the simulation data to the final 400 ns for analysis, the root-mean-square fluctuations (RMSF) were calculated for the Cα atoms of the histones and the average of the nucleic acids for DNA. The RMSF is used to calculate local time-averaged fluctuations. The RMSF of DNA (**Fig S1**) was calculated for thirds of the final 400 ns and then the standard deviation of the mean plotted. Contact analysis was calculated with a cutoff distance of 8 Å between histone Cα atoms to compare dimer interface contacts in both systems. The center-of-mass (COM) of dimers was then calculated along the trajectory and the distribution of distances between COMs compared.

Principle component analysis (PCA) was performed on the histone core as previously described^2^. This analysis was then extended to include Cα atoms of histones and phosphate atoms nucleosomal DNA. The first and last ten base pairs were truncated from the analysis to remove ends calculated to have a high RMSF (**Fig S1**). This alteration was made so that DNA end motions did not dominate the major principle components. The magnitude of motion is multiplied by a factor of 5 in the movies to amplify motions for visual clarity.

### Cloning

GFP-CENP-A plasmids were a gift from Stephan Diekmann. To mutate K124 to alanine (A) or glutamine (Q) residues, fusion PCR was performed using a reverse primer (TGGGAAGAGAGTAACTCGG) along with a forward primer from the 5’ START codon that includes an EcoRI site. That amplicon was gel purified and combined with a PCR amplicon that used a forward primer encompassing the K to [A] or [Q] mutation (CGAGTTACTCTCTTCCCA[GCG]GATG or CGAGTTACTCTCTTCCCA[CAG]GATG, respectively) and a reverse primer that includes the XmaI site and STOP codon. The final fusion PCR product was excised, gel purified and finally ligated downstream of plasmid that had either GFP or HA-tags, driven by a constitutive CMV promotor.

### Transfections

HeLa cells were grown to ˜75% confluency and transfected using Lonza’s Amaxa Cell Line Nucleofector Kit R (Cat #VCA-1001) using Amaxa Biosystems Nucleofector II electroporation system according to the manufacturer’s guidelines using program O-005. After transfection, cells were plated with fresh media and grown for 48 hrs before harvesting for experiments.

### Cell synchronization, native chromatin immuno-precipitation (nChIP)

HeLa cells were grown in DMEM (Invitrogen/ThermoFisher Cat #11965) supplemented with 10% FBS and 1X Pen/Strep. nChIP experiments were performed without fixation, and with or without a double thymidine block to synchronize cells. For the complete double thymidine block protocol, please refer to Bui *et al*, 2012.

After cells were harvested, they were washed with PBS and PBS containing 0.1% Tween 20. Nuclei were released with TM2 (20 mM Tris-HCl, pH: 8.0; 2 mM MgCl_2_) with 0.5% Nonidet P 40 Substitute (Sigma Cat #74385). Afterwards, nuclei were washed with TM2 and chromatin were either digested for 4 min for nChIP or 8 min for ChIP-seq with 1.0 U MNase (Sigma Cat #N3755-500UN) in nuclei solubilized with 2 mL of 0.1 M TE (10 mM Tris, 0.2 mM EDTA, 100 mM NaCl) and supplemented with 1.5 mM CaCl_2_. MNase reactions were quenched with 10 mM EGTA and centrifuged at 1000 rpm at 4°C. Supernatant was removed and nuclei extracted over-night at 4°C in 0.5X PBS supplemented with a protease inhibitor cocktail (Roche Cat #05056489001). ChIP was performed with anti-HA antibody (Santa Cruz Cat #sc-805). nChIP’ed chromatin bound to Protein G Sepharose beads (GE Healthcare Cat #17-0618-02) were washed 3X with cold 0.5X PBS. Westerns were done using LiCor’s Odyssey CLx scanner and Image Studio Ver 2.0. Antibodies used for Westerns include: CENP-A (AbCam Cat #ab13939), CENP-B (AbCam Cat #ab25734, CENP-C (MBL Cat #PD030), HA-tag (GenScript Cat #A01244), FLAG-tag (AbCam Cat #ab1162) and H2B (AbCam Cat#52484).

### TAU gels and Westerns

For synthesis, prepatory and running conditions for TAU gels, please refer to Walkiewicz *et al*. (2014). For Western transfers, we used the Trans-blot Turbo Transfer Pack (mini Biorad Cat #170-4158 or midi Biorad Cat #170-4159). For Western detection of the proteins, we used LiCor’s secondary infrared antibodies, the Odyssey CLx laser scanning system and Image Studio Ver 2.0 to quantify the protein levels.

### Immuno-fluorescence (IF)

For complete cell and chromatin fiber IF protocols, please refer to Bui *et al*, 2012. GFP tags were fused in-frame and upstream of wild-type CENP-A, K124A and K124Q, and may be co-transfected with mCh-CENP-A for co-localization assays. IF was performed using CENP-B (AbCam Cat #ab25734) and CENP-C (MBL Cat #PD030) antibodies. Cells were pulsed with EdU 30 min prior to the desired time-point using the Click-iT EdU Alexa Fluor 594 kit (Life Technologies Cat #C10639), and imaged using a DeltaVision RT system fitted with a CoolSnap charged-coupled device camera and mounted on an Olympus IX70. Deconvolved IF images were processed using Image J and to assess colocalization with its ‘Colocalization Finder’ plug-in.

### LC-MS

Tau gel bands were processed using an in-gel digestion protocol from Shevchenko et al *(Nature Protocols* **1**, 2856 – 2860 (2007)). Each band was split in two and separate trypsin and chymotrypsin in-gel digestions were performed. The samples then underwent shotgun proteomic analysis on a nano-HPLC system (NanoLC 2D; Eksigent, Dublin, CA) coupled to a hybrid mass spectrometer (Orbitrap Velos Pro; Thermo-Electron, Bremen, Germany). Samples were injected using an auto-sampler and loaded onto a self-packed trap column (2cm, 100μm ID, packed with C18 Magic AQ from Michrom Bioresources, Auburn, CA), and the samples were then analyzed on a self-packed C18 (15cm, 2.7μm HALO Peptide ES C-18, MAC-MOD, Chadds Ford, PA) column with a laser pulled tip (P-2000, Sutter, Novato, CA) using a flow rate of 200nL/min. The column was heated to 50°C using column heater (Phoenix S&T, Chester, PA). Mobile phase A was water with 0.1% formic acid and mobile phase B was acetonitrile with 0.1% formic acid. The analytical gradient was a 90 minute linear gradient from 5 to 35% buffer B. Eluting peptides were electrosprayed at 2.3kV, and the ion transfer capillary was heated to 250°C. The Orbitrap was operated in data-dependent mode with different settings depending on the cleavage enzyme used: Trypsin cleaved samples were analyzed with a CID top 18 method, and chymotrypsin cleaved samples were analyzed with a CID and ETD decision tree top 12 method. Precursor resolution was set to 60,000, CID collision energy was 35%, and ETD time reaction time was 100ms with supplemental activation.

### Database Search Parameters

Protein identification was performed against the UniProt database entry for CENPA using Proteome Discoverer 2.1 (Thermo Fisher Scientific) equipped with SEQUEST HT and Mascot (Matrix Science, Boston, MA). Search settings included tryptic or chymotryptic digest with up to 2 missed cleavages or non-specific cleavage. Carbamidomethylation of cysteine was set as a static modification while dynamic modifications included Met oxidation, Asp, Glu deamidation, Ser, Thr, Tyr phosphorylation, Lys acetylation, Lys ubiquitination, Arg and Lys methylation. Only matches with XCorrs greater than 2.0 or ion scores greater than 20 were considered. All the spectra matches were manually validated.

## ACKNOWLEDGEMENTS

We thank Drs. Tom Misteli, Sam John, Mirit Aladjem and members of the CSEM lab, particularly Daniel Melters for critical comments and suggestions.

## AUTHOR CONTRIBUTIONS

MD and YD designed the biological study; MP, YD and GP designed the computational study; MB performed all biochemical and cell biology experiments with assistance from PDA and SR; MP performed all computational experiments; MP, YD and GP analyzed the computational experiments; AN and ANL performed and analyzed the MS/MS experiments; MB, MP, GP and YD wrote the manuscript.

## Supplementary Figure & Movie Legends

**Figure S1.**
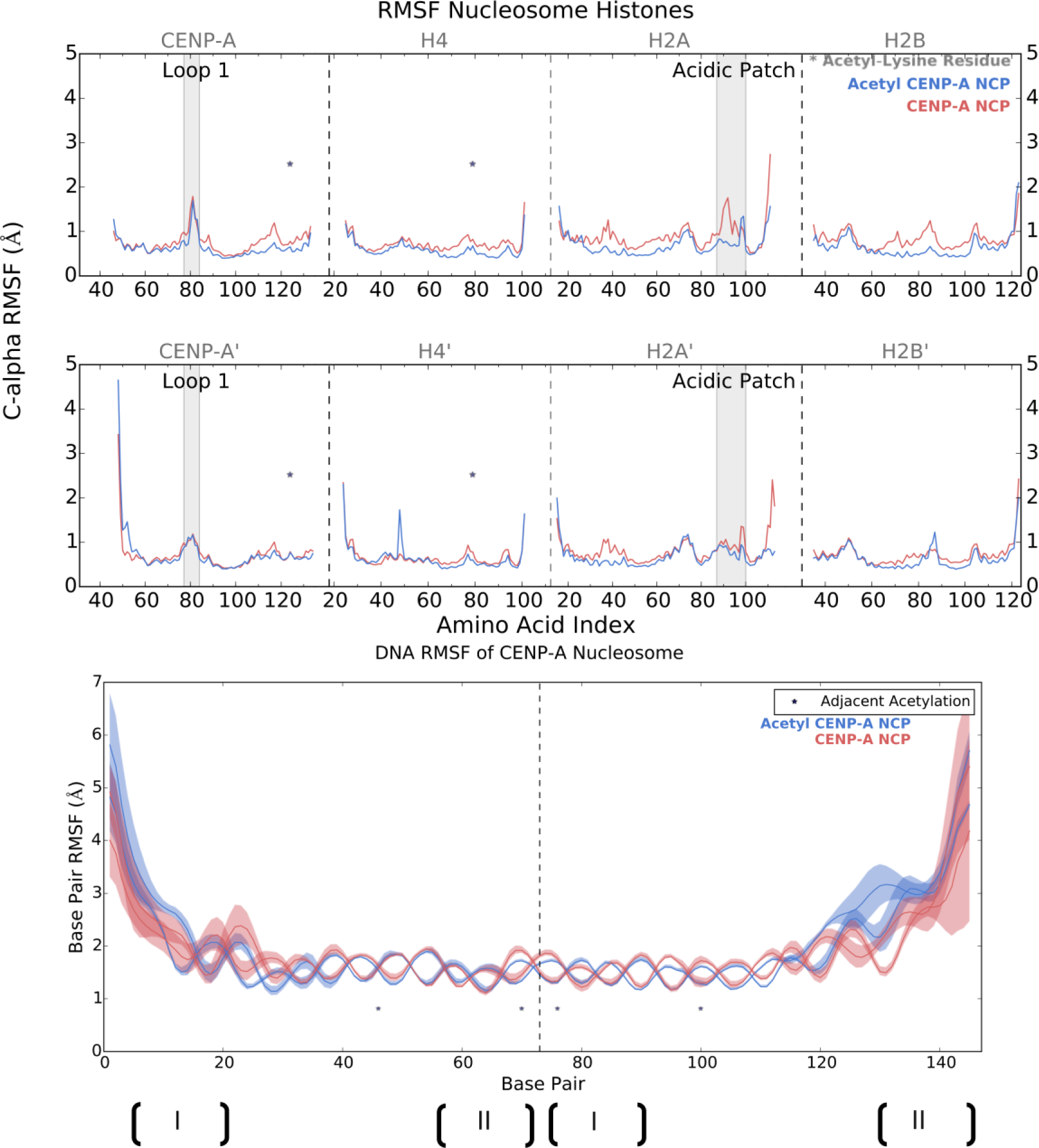
RMSF of proteins. **A)** This decrease in RMSF of Cα residues upon acetylation is more pronounced on the histone heterotetramer adjacent to the entry DNA. Of particular interest, the RMSF of the acetylated H2A acidic patch was suppressed with acetylation by −1 Å and suppression is shown in the CENP-A C-terminus. The greater similarity shown in the RMSF of the reciprocal histones—CENP-A’, H4’. H2A’, and H2B’—could potentially be explained by the observed asymmetric unwrapping of DNA where the exit end in both system dissociates to a similar amount (**Fig 2A**). **B)** The RMSF of whole base-pairs is shown for each DNA strand. Regions marked by I are DNA wrapped near the entry or near CENP-A and II are near the exit end of CENP-A’. The pseudo-dyad is marked by the vertical dotted line.

**Figure S2.**
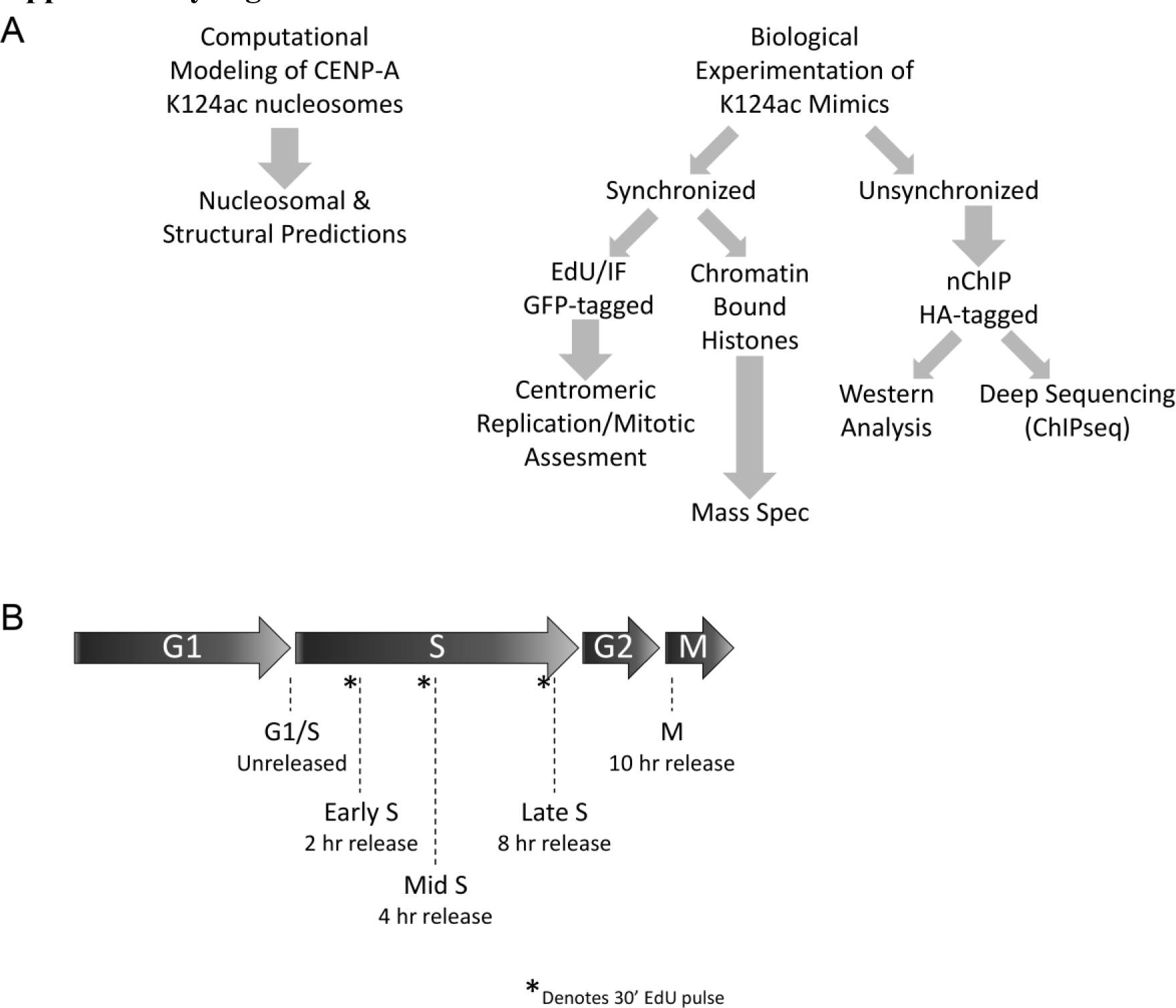
Experimental scheme for experiments and cell synchronization. **A)** Computation and biological experimental scheme for this publication. **B)** Cell cycle synchronization with a 30 min EdU pulse prior to preparing slides for EdU and staining immuno-fluorescence.

**Figure S3.**
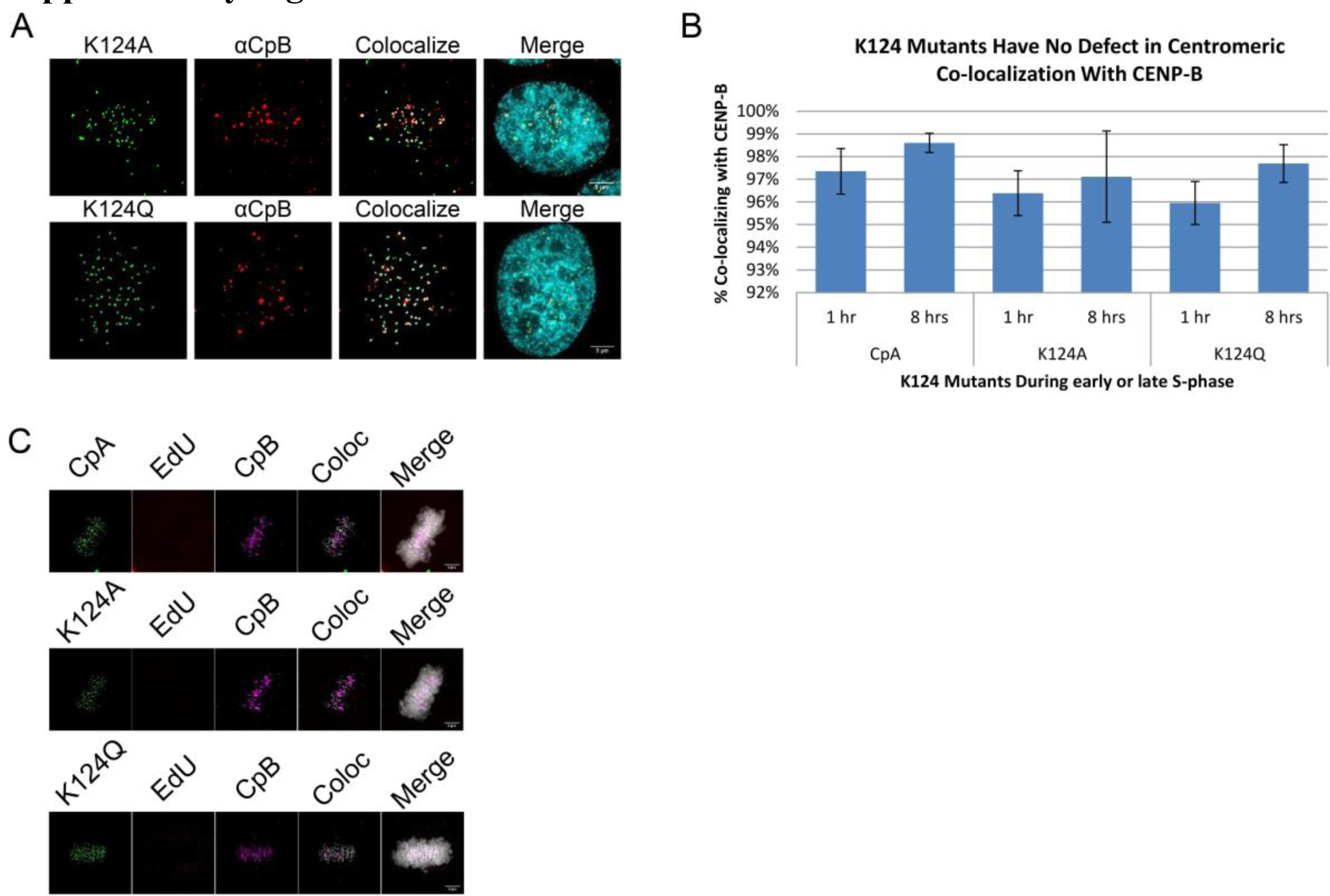
Co-CENP-B staining with HA-tagged K124A/Q proteins. **A)** Unsynchonized cells stainined with CENP-B (CpB) and **B)** percentage of CENP-B colocalizing with GFP-CENP-A (CpA) or K124A/Q exogenously expressed proteins after a double thymidine block and released for 1 or 8 hrs. **C)** Colocalization of the GFP-CpA or K124A/Q with CpB during metaphase or mitosis.

**Figure S4.**
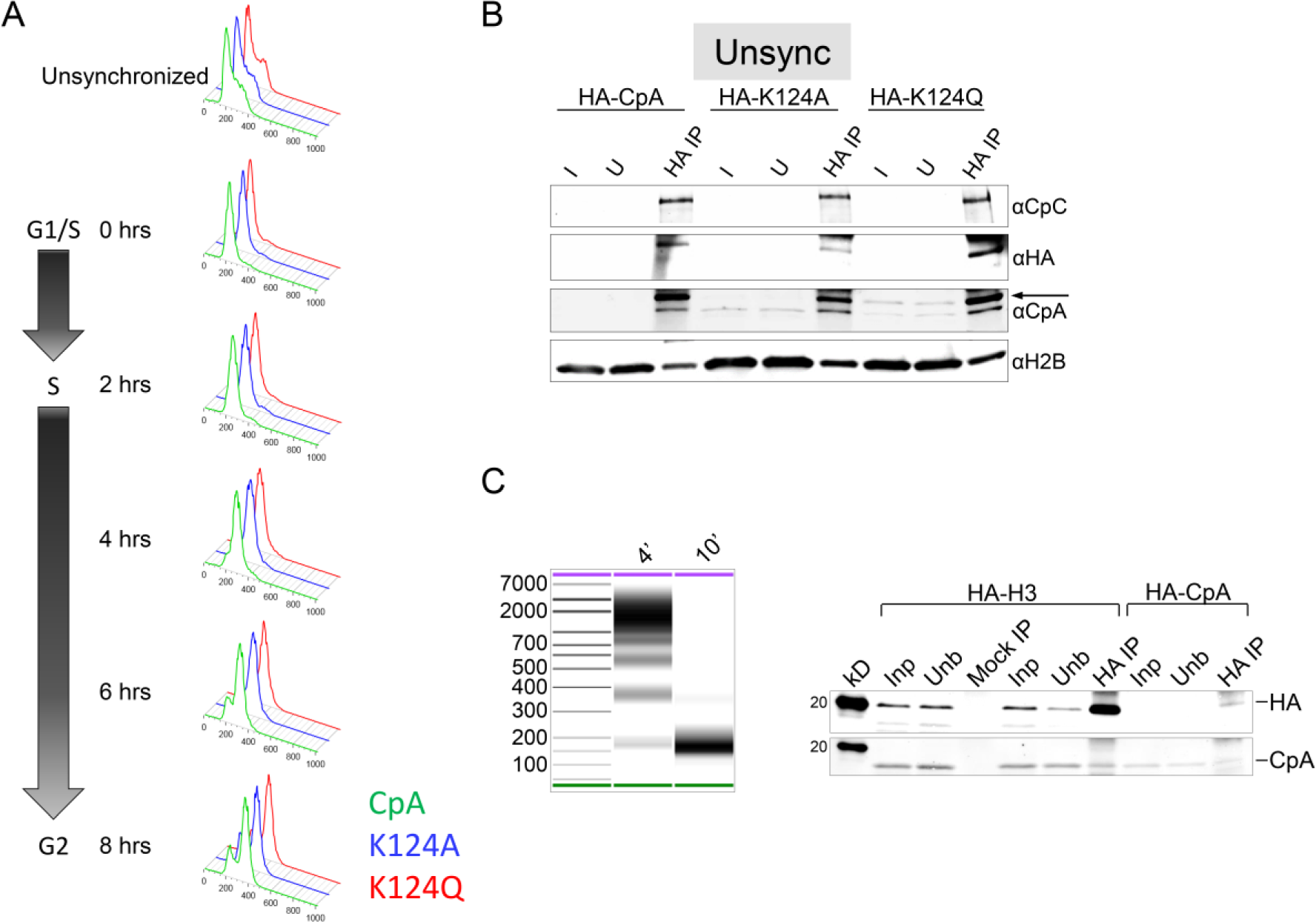
K124A/Q mutants have normal cell ploidy but altered CENP-C affinity. **A)** FACS analysis of GFP-CpA and K124A/Q mutants. X-axis is the Propidium Iodide level and the Y-axis is the cell count. **B)** HA-tagged proteins were ChIP’ed from HA-CpA and K124A/Q mutants and probed by Western against CENP-C (CpC), HA, CpA and H2B. **C)** Two-week old geneticin selected stable cells expressing either HA-H3 or HA-CpA with medium length DNA arrays(4’ MNase) (versus very short arrays (10’ MNase) on BioAnalyzer) were ChIP’ed using anti-HA antibodies. Unlike HA-H3, HA-CpA stable cells are actively repressing exogenous CENP-A protein production/incorporation. kD mark =20kD, I=Input, U=Unbound.

**Figure S5.**
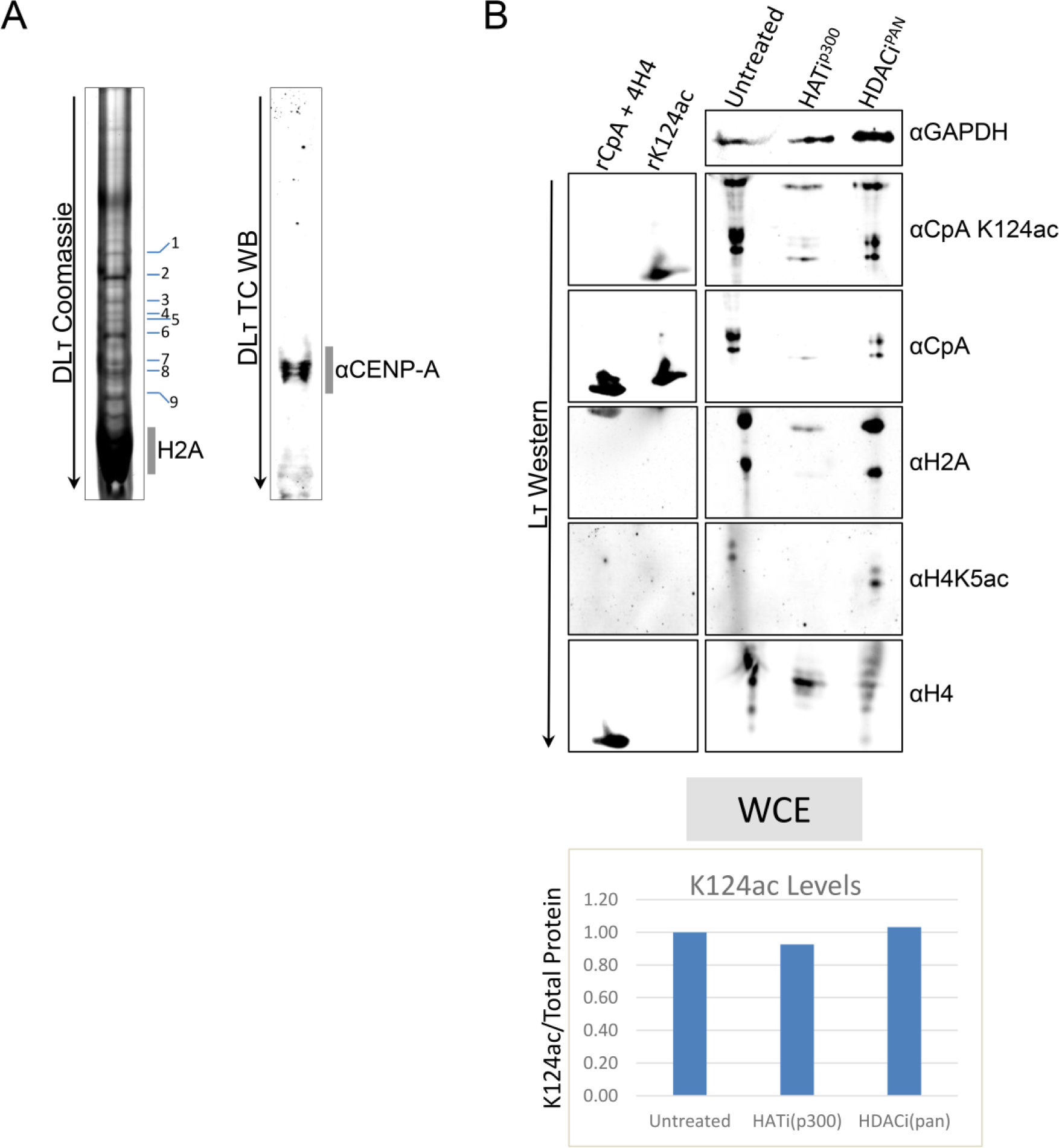
Identifying CENP-A on Double Long TAU (DLτ) and Long TAU (Lτ) gels for mass spectrometry and HAT/HDAC inhibitor drug treatments. A) dLτ gels remove excess canonical histone components, leaving behind predominantly CENP-A and histone H2A. Numbers on the DLτ represents bands that were sent for mass spec and duplicate gel was used for Western and probed against CENP-A. B) Whole cell extracts from cells expressing HA-CENP-A (HA-CpA), HA-K124A and HA-K124Q were resolved on SDS-PAGE gels and analyzed on Western blots against CENP-C, HJURP and CENP-A. Black arrow indicates exogenously transfected HA-tagged proteins. Lowest kD marker (Invitrogen Magic Mark XP Cat #LC5602) is 20 kD.

### Movie S1 (histones only, pca_histone_ms.mov)

Histone dimers CENP-A/H4 is shown in red, CENP-A’/H4’ in blue, H2A/H2B in light blue, and H2A’/H2B’ in white. Movies presented here are animations of the most significant mode of motion, PC1^core^, of the Principal Component Analysis of histones (PCA^core^). The first clip shows the histones in both systems rocking with a “freezing” of motion in the acetyl CENP-A histones on the right. The second clip shows the 4-helix bundle in isolation to highlight the interface formed between CENP-A and CENP-A’. Next, the histones are rotated to focus on the described scissoring motion between helices α2 and α3 in the acetyl NCP. Here, the two helices move apart and then together, modulating the widths of the major and minor grooves in the acetyl NCP. Next, flipped to the other side of the nucleosome core, observe the rigidification of the H2A/H2B to H2A’/H2B’ interface in the acetylated nucleosome. To clarify this further, we then show the H2A to H2A’ interface in isolation.

### Movie S2 (whole nucleosome, pca_movies_ms.mov)

Histone dimers CENP-A/H4 is shown in red, CENP-A’/H4’ in blue, H2A/H2B in light blue, and H2A’/H2B’ in white. Presented are animations of the most significant mode of motion of the whole nucleosome, PC1^nuc^, of the Principal Component Analysis (PCA^nuc^). The pseudo-dyad is labeled PD and the modified lysine side chains are shown in green with or without acetylation dependent on the system. It is worth noting that our PCA^nuc^ calculations are based on DNA phosphate positions and protein Cα’s—therefore, side chains are stagnant relative to the protein backbone. In the first two clips, two unique features of the acetyl NCP are shown: the modulation in the width of the major and minor DNA grooves and the inter-helical DNA bubble formed adjacent to H4 and H4’ K79ac. The final clip then shows the NCP on the side to highlight DNA end untwisting in the acetyl NCP with the last ten base pairs were truncated from the analysis.

